# A trajectory-coupled network bottleneck governs gemcitabine resistance in 3D PDAC tissue models

**DOI:** 10.64898/2026.03.24.713885

**Authors:** Maryam Almasi, Jesús Guillermo Nieves Pereira, Thomas Dandekar, Gudrun Dandekar, Johannes Balkenhol

**Author notes:** These authors contributed equally to this work as first authors (Maryam Almasi, Jesús Guillermo Nieves Pereira). These authors contributed equally to this work as senior authors (Thomas Dandekar, Gudrun Dandekar, Johannes Balkenhol). Correspondence: Johannes Balkenhol.

## Abstract

PDAC exhibits rapid chemoresistance, yet how drug-tolerant states arise remains unclear. Existing approaches miss how network topology evolves across cell-state transitions under drug pressure.

A 3D PANC-1 tissue model on decellularized intestinal matrix was used for scRNA-seq across four conditions (control, GEM, TGF-β1, GEM+TGF-β1). Pseudotime trajectory inference was combined with dynamic PPI network analysis. Findings were cross-examined in a PDAC atlas (726,107 cells, 231 patients; Loveless et al., 2025).

GEM resistance involved E2F1, mTOR, CDK1, AURKA, TPX2, TOP2A, and BIRC5. TGF-β1 drove EMT resistance via KRAS, glycolysis, and hypoxia, inducing SPOCK1, MBOAT2, COL5A1, ADAMTS6, THBS1, and FN1. Trajectory-coupled network analysis revealed an emergent bottleneck when G1→S and TGF-β1-induced EMT co-occurred: CDK1 centrality spiked selectively, with CDKN1A as critical regulator. This CDK1–CDKN1A–WEE1 axis defines an “S-phase persistence” state enriched for GEM survivors. Atlas cross-examination confirmed 8.7-fold metastatic enrichment of triple-positive cells and EMT–cell-cycle coupling.

Trajectory-coupled network topology analysis identifies CDK1–CDKN1A–WEE1 as a chemoresistance bottleneck corroborated in 726,107 patient cells. The framework generalizes to drug resistance across cancer types.

## Introduction

Pancreatic ductal adenocarcinoma (PDAC) is a disease with high chemoresistance and mortality (Siegel et al., 2024). Late diagnosis hinders curative treatment success after surgery. Standard chemotherapeutic treatments include Gemcitabine (GEM) combined with fluoropyrimidine-based treatment regimens (Sohal et al., 2018; Hosein et al., 2022), resulting in still poor median overall survival (OS) rates of approximately one year (Conroy et al., 2011; Hosein et al., 2022). A more recent clinical study shows better efficacy of adjuvant FOLFIRINOX (oxaliplatin, irinotecan, leucovorin, fluorouracil) than GEM alone (Conroy et al., 2022), though with a less favorable safety profile (Conroy et al., 2018).

GEM is a nucleoside analog used as a chemotherapeutic agent for over 25 years. A major problem is poor tumor penetration due to high stroma content and lack of vascularization, particularly in PDAC (de Sousa Cavalcante et al., 2014; Neesse et al., 2011). As a prodrug, GEM must be metabolized to its active form and cellular uptake is mediated by human nucleoside transporters (hNTs) — all of which can mediate resistance (de Sousa Cavalcante et al., 2014). Critically, although GEM effectively blocks DNA synthesis, subsequent cell-cycle checkpoints remain delayed even after drug removal (Cappella et al., 2001), suggesting that dynamic cell-state transitions under drug pressure — rather than static expression differences alone — govern which cells survive.

This emphasizes the urgent need for physiologically relevant preclinical models that faithfully recapitulate in vivo chemoresistance and enable mechanistic dissection of drug-tolerant cell states. We address this by utilizing a 3D intestinal tissue matrix-based tumor model (SISmuc: Small Intestinal Submucosa with preserved mucosa) harboring the PANC-1 cell line, which carries common PDAC driver mutations including KRAS, TP53, and CDKN2A (Deer et al., 2010). Unlike conventional 2D cultures with artificially high proliferation, our 3D model reaches a homeostatic state after ∼11 days with proliferation indices (∼35% Ki67-positive) closely matching the clinical mean in PDAC patients (Striefler et al., 2016; Göttlich et al., 2018; Peindl et al., 2022), validating it as a suitable platform for studying chemoresistance under near-physiological conditions.

To investigate resistance mechanisms, we utilized this matrix, which preserves extracellular matrix (ECM) components and provides distinct tissue niches within former crypt and villus structures, thereby enabling tissue-niche–specific responses to GEM treatment. To mimic the desmoplastic reaction characteristic of PDAC (Neesse et al., 2011), we further stimulated PANC-1 tissue models with TGF-β1 and compared GEM responses with and without stimulation. The resulting single-cell dataset was generated from PANC-1 tissue models prepared in our laboratory, with sequencing performed at a dedicated facility.

This dataset has been explored in two complementary directions: a mathematical strategy approach using pseudo-bulk RNA-seq identified genes upregulated in GEM-surviving versus untreated cells (Caliskan et al., 2025). Here, we complement this work at single-cell resolution by integrating cell-state trajectory inference with dynamic regulatory network analysis — tracking how network topology changes as cells transition between drug-exposed states — to resolve the mechanistic decision points that drive GEM resistance rather than merely marking it. We examine the molecular composition of distinct PANC-1 cancer subgroups in detail, focusing on their cell-cycle progression and the impact of TGF-β1 stimulation.

There is evidence that EMT and DNA damage response (DDR) components are reciprocal regulators of TGF-β1 signaling and cell-cycle progression (Schuhwerk et al., 2023), and that cell fate is governed by a limited set of core regulatory circuits reaching internal equilibrium during tumor progression (Lee et al., 2024). ScRNA-seq enables identification of vulnerabilities in cancer subpopulations exposed to chemotherapeutics precisely because DDR plasticity increases genetic variability when the cell cycle is not fully arrested — making dynamic, single-cell resolution essential to capture the transitional states that govern resistance.

This dataset has been explored in two complementary directions: a mathematical strategy approach using pseudo-bulk RNA-seq identified genes upregulated in GEM-surviving versus untreated cells (Caliskan et al., 2025). Here, we complement this work at single-cell resolution by integrating cell-state trajectory inference with dynamic regulatory network analysis — tracking how network topology changes as cells transition between drug-exposed states — to resolve the mechanistic decision points that drive GEM resistance rather than merely marking it. We examine the molecular composition of distinct PANC-1 cancer subgroups in detail, focusing on their cell-cycle progression and the impact of TGF-β1 stimulation.

To pinpoint these decision points, we reconstructed regulatory networks at single-cell resolution and monitored their topology changes along cell-state trajectories — resolving intermediate and transitional states that binary treated-versus-untreated comparisons would miss. This approach identified major control nodes, along with their upstream and downstream regulators, acting as high-impact bottlenecks governing cell fate under drug pressure. To assess whether this regulatory architecture reflects human tumor biology, we cross-examined key findings in a comprehensive PDAC single-cell atlas comprising 726,107 cells from 231 patients across 12 independent studies (Loveless et al., 2025). The resulting framework not only identifies marker genes of resistance but highlights intervention points directly relevant to ATR–CHK1–WEE1 and TGF-β1/SMAD3 signaling axes currently being targeted in clinical trials for PDAC.

## Material and Methods

### Cell culture

PANC-1 cells (purchased from DSMZ, number ACC-783) were maintained in Dulbecco’s Modified Eagle Medium (DMEM, high glucose, GlutaMAX™; Gibco, 61965059, Germany) supplemented with sodium pyruvate (Invitrogen, P2256, Germany) and 10% fetal calf serum (FCS; PAN-Biotech, P303306, Germany). Cells were cultured in T-75 or T-150 flasks (Techno Plastic Products, TPP 90076/90151, Switzerland) at 37 °C and 5% CO₂ and passaged at 80–90% confluence.

For passaging, 3–6 mL of 1% trypsin prepared in PBS without calcium and magnesium (PBS⁻; Sigma-Aldrich, D8537-500ML, Germany) supplemented with 0.5 mM ethylenediaminetetraacetic acid (EDTA; Sigma-Aldrich, E5134, Germany) was added to T-75 or T-150 flasks, respectively, and incubated for 3 min. Detachment was quenched by adding either twice the volume of FCS-supplemented culture medium or 2 mL of FCS. Cell suspensions were transferred to 50 mL tubes (Greiner Bio-One, 188271N, Germany) and centrifuged at 300 g for 5 min (Thermo Fisher Scientific, Multifuge X12, Germany). Supernatants were removed with a disposable Pasteur pipette connected to a vacuum pump (KNF Neuberger, Germany), and pellets were gently resuspended to the desired volume in complete medium using 5- or 10-mL serological pipettes (Greiner Bio-One, 606180/607180, Germany).

### Generation and culture of 3D SISmuc models

Explanation of porcine jejunum were in compliance with the German Animal Protection Laws (§4 Abs. 3) and all animals received humane care in compliance with the guidelines by the FELASA, WHO and FDA (WHO-TRS978 Annex3 and FDA-OCTGT Preclinical Guidance) after approval from the institutional animal protection board (registration reference number#2532-2-12, Ethics Committee of the District of Unterfranken, Würzburg, Germany). Decellularization was performed as previously published (Linke et al, 2007). In short, the intestinal porcine segments were extensively rinsed with PBS buffer, then chemically decellularized with a sodium deoxycholate monohydrate solution, rinsed again, and subsequently underwent gamma sterilization. All steps for 3D tissue model generation were performed in a sterile biosafety cabinet. SISmuc scaffolds were incised with a scalpel (Bayha handle 504 with blade N22, Germany) and sectioned into ∼150 × 150 mm squares. Using sterile forceps (Bochem Laborbedarf, 1023, Germany), each piece was mounted between two sterile ring-like inserts (“cell crowns”) and transferred to a sterile 12-well plate (TPP, 92012, Switzerland). To prevent dehydration, 2.5 mL of complete medium was added per well in a transwell-like configuration: 1.5 mL to the outer compartment (between the well wall and crown) and 1.0 mL to the inner compartment. The luminal surface of the former intestinal tissue was oriented upward (inner compartment).

PANC-1 SISmuc 3D models were established by seeding 2 × 10⁵ cells/mL; 500 µL of the cell suspension was carefully dispensed onto the luminal surface of each construct. After a 1-hour attachment period, 1 mL of complete medium was added to both the luminal and basolateral compartments.

For routine medium exchange, spent medium was removed from both compartments with disposable glass Pasteur pipettes and replaced with 1.0 mL (inner) and 1.5 mL (outer) of fresh medium at each scheduled change, unless otherwise stated.

### Stimulation and drug treatment of 3D models

Gemcitabine (GEM; Sigma-Aldrich, G6423-10MG, Germany) was dissolved in sterile DMEM to a concentration of 10 mM and then diluted in complete medium to 10 µM. It was added directly to both compartments in the volumes previously described. GEM treatment was applied after 11 days of culture and applied for 24h.

For transforming growth factor beta 1 (TGF-β1) stimulation, recombinant human TGF-β1 (Peprotech, 100-21-1006, Germany) was reconstituted in PBS containing 1% bovine serum albumin (BSA; Carl Roth, 01633, Germany) to 10 µg/mL and diluted to 10 ng/mL in complete medium immediately before use, and added directly to both compartments. TGF-β1 stimulation was constantly kept between days 3-14 of each TGF-β1+ model with refreshing during each medium change.

### Hematoxylin–eosin and immunofluorescence staining

SISmuc constructs were fixed, sectioned, deparaffinized, and rehydrated as described previously (any previous ref of SISmuc handling).

For HE (Hematoxylin–eosin) stainings, pre-treated sections were immersed in hematoxylin (Morphisto GmbH, 61870036, Germany) for 6 min, rinsed in deionized water, and blued under running tap water for 5 min. Sections were then immersed in eosin (Morphisto GmbH, 10177.01000, Germany) for 6 min and rinsed with deionized water. Dehydration/clearing was performed by a brief dip in 70% ethanol (prepared from absolute ethanol (Carl Roth, K928.4, Germany) diluted with deionized water), followed by 2 min in 96% ethanol, 5 min in Isopropanol (Carl Roth, T910, Germany), and 10 min in xylene (Carl Roth, 9713, Germany). Slides were mounted with Entellan® (Merck, 1079600500, Germany) and allowed to dry overnight prior to imaging.

On the other hand immunofluorescent staining with Ki67 antibody (Abcam, ab1666, Germany) was subjected to heat-mediated antigen retrieval by incubation for 20 min at 100°C in citrate buffer (pH 6.0; 42 g/L citric acid monohydrate (Carl Roth, 1002441000, Germany) adjusted with NaOH) using a steam cooker (Braun, FS20, Germany), then rinsed in deionized water and allow to cool down to RT. Thereafter, hydrophobic barriers were drawn with ImmEdge® Hydrophobic Pen (Vector Laboratories, H-4000, USA). Sections were transferred to PBS-T (PBS containing 0.5% Tween-20 (Sigma-Aldrich, P9416, Germany)).

Sections were blocked for 20 min at room temperature with 5% donkey serum (Sigma-Aldrich, D9663, Germany) in antibody diluent (DCS Innovative Diagnostic Systems, AL120R500, Germany). The primary antibody was diluted 1:100 in antibody diluent and incubated overnight at 4 °C. Negative controls received diluent only.

After three 5-minute washes in PBS-T, sections were incubated for 1 hour at room temperature, protected from light, with secondary antibodies diluted 1:400 in antibody diluent: anti-rabbit Alexa Fluor 555 (Life Technologies, A-31572, USA). Following three additional washes in PBS-T in the dark, slides were mounted with Fluoromount-G™ with DAPI (Invitrogen, SBA-0100-20, USA) and allowed to dry overnight prior to imaging.

### MTT Assay

In 3D matrix-based models, viability was determined by an MTT assay. For this, 1 mg/mL MTT solution (SERVA, Heidelberg, Germany) was added to the samples and incubated for 3 hours under standard conditions. Afterwards, the medium was removed and the 3D matrix model was transferred to an empty 50 mL centrifuge tube. Thereafter, formazan was washed out with 2 mL of Isopropanol with 0.04 N HCl and kept in an orbital shaker for one hour. Then the supernatant was collected in a fresh 15 mL centrifuge tube, and an extra milliliter of Isopropanol HCl was added. This was kept in the same conditions for 30 minutes and repeated once. After mixing, 200 μL of each sample was pipetted into a 96-well microplate, and the absorbance at 570 nm was measured using a microplate reader (Tecan, Männedorf, Switzerland).

### Preparation of single-cell suspensions for scRNA-seq

Media was removed from both compartments, and constructs were washed three times with room-temperature PBS⁻. Pre-warmed Accutase® (1 mL; Sigma-Aldrich, A6964, Germany) was added to the luminal side and incubated for 30 min with gentle pipette trituration every 10 min; the suspension was collected into a 15 mL tube. Constructs were then incubated for 10 min at room temperature with 10 mg/mL protease (Bacillus licheniformis subtilisin A; Sigma-Aldrich, P5380, Germany), mixing at 5-minute intervals; the suspension was collected, and the crowns were rinsed twice with 1 mL PBS⁻. All fractions were pooled, the volume was adjusted to 5 mL, and cells were centrifuged at 300 g for 5 min. Pellets were resuspended for counting and subjected to dead-cell removal using the manufacturer’s protocol (Miltenyi Biotec, 130-090-101, Germany). Cell concentration was adjusted for optimal capture, and samples were processed according to 10x Genomics tagging and library preparation guidelines (10x Genomics).

### Single-cell RNA sequencing and processing

Single-cell sequencing was performed at the Helmholtz Institute for RNA-based Infection Research (HIRI, Single-Cell Center) using the 10x Genomics Single Cell 3′ v3 chemistry. Libraries were processed with Cell Ranger v7.0.1 (10x Genomics) for demultiplexing, alignment, and UMI counting, including intronic reads. Technical parameters of eight tagged samples are shown in Table S1.

### Preprocessing, quality control, and clustering

Raw FASTQ files were processed with Cell Ranger v9.0.0 (10x Genomics) against GRCh38 (RefSeq, October 2023) using the *multi* pipeline for demultiplexing/quantification and the *aggregate* command for merging per-sample outputs (10x Genomics, n.d.). The run configuration (gene-expression reference, library specs, Cell Multiplexing Oligos mapping) was defined in cellranger_config.csv. Aggregated matrices and metadata were imported to AnnData for downstream analysis.

Cell Ranger outputs (barcodes.tsv, features.tsv, matrix.mtx) were parsed with Pandas and SciPy (McKinney, 2010; Virtanen et al., 2020) and assembled into per-sample AnnData objects, which were concatenated with Scanpy (Wolf et al., 2018) while preserving unique cell/gene identifiers. Gene symbols were mapped from Ensembl IDs via MyGene (Xin et al., 2016). Features labeled “CMO” were removed.

Quality control (QC) thresholds were: mtRNA < 10%, rRNA < 50% (MT-, RPS-/RPL-prefixes), n_genes 1,900–9,000, and UMIs 1–70,000; cells outside these bounds were discarded. Scrublet was used for doublet filtering (Wolock et al., 2019). Extreme outliers were removed using a 5× MAD rule on log-transformed counts. Counts were library-size normalized to 1×10⁴ UMIs per cell and log1p-transformed. Highly variable genes (HVGs) were selected with default Scanpy settings (top 2,000–4,000 by dispersion/mean). Processed datasets were saved in H5AD format for reproducibility.

Dimensionality reduction used PCA (ARPACK solver), followed by a k-nearest-neighbor graph on the first 40 PCs with k = 8 (Wolf et al., 2018). UMAP was computed for visualization (McInnes et al., 2018), and unsupervised clustering used the Leiden algorithm (Traag et al., 2019). Clustering resolution was swept from 0–1 in 0.01 steps to obtain stable partitions; an outlier cluster was removed and the graph/UMAP recomputed, yielding four robust clusters. Plots were generated with matplotlib and seaborn (Hunter, 2007).

Reproducibility: Exact commands, configuration CSVs, random seeds, intermediate file names (e.g., H5ADs), and figure-generation code are provided in the accompanying Jupyter notebooks (see github link provided).

### Differential expression (DE)

Cells were grouped by Leiden cluster × treatment (e.g., *Leiden3_GEM2* vs *Leiden0_GEM2*) from the prefiltered AnnData (H5AD). DE was performed with Scanpy rank_genes_groups using a t-test; we report log₂ fold change, raw *p*, and BH-FDR–adjusted *p* values (Wolf et al., 2018). Summary tables (including basemean per gene) and figures (volcano/bar plots) were generated in Python using Pandas, SciPy, matplotlib, and seaborn (McKinney, 2010; Virtanen et al., 2020; Hunter, 2007; Waskom, 2021).

### Gene set enrichment analysis (GSEA)

Ensembl-IDs were mapped to symbols via MyGene.info (Wu et al., 2013). GSEA was run with GSEApy (Fang et al., 2022) on MSigDB Hallmark 2020 gene sets with 1,000 permutations, min size = 10, max size = 50, and the log₂ ratio of classes scoring metric. We report NES, *p*, FDR, and leading-edge genes. Optional summaries included enrichment maps drawn with NetworkX and pathway lookups via Reactome2py (Hagberg et al., 2008; Jassal et al., 2020).

### Enrichment of GEM-treated cells within Leiden clusters

GEM treatment status (metadata field GEM) was summarized per Leiden cluster as % GEM cells and visualized on UMAP using Scanpy (Wolf et al., 2018). Statistical enrichment was tested per cluster using Fisher’s exact test (SciPy stats.fisher_exact), comparing in-cluster vs out-of-cluster counts of GEM vs non-GEM cells; *p* values were BH-FDR–adjusted across clusters (Virtanen et al., 2020). Results (odds ratios, adjusted *p*) were compiled into a summary DataFrame (Pandas) and exported.

### Cell-state and fate inference (trajectory analysis)

We analyzed QC’d, HVG-filtered AnnData using scFates v1.0.6 (Faure et al., 2023), Palantir v1.3.3 (Setty et al., 2019), and Scanpy (Wolf et al., 2018) in Python 3.8 with Pandas, NumPy, SciPy, rpy2, and Matplotlib (McKinney, 2010; Harris et al., 2020; Virtanen et al., 2020; Hunter, 2007).

PCA embeddings were passed to Palantir to compute diffusion maps and a multiscale diffusion space (n_eigs=4). This representation was stored as X_palantir, a new neighbor graph was computed on it, and a ForceAtlas2 embedding was initialized from X_pca2d for visualization.

Trajectory trees were inferred with scFates using the Principal Tree (PPT) method on the Palantir space, exploring multiple settings; one representative configuration was Nodes=100, seed=42, ppt_lambda=1, ppt_sigma=0.08, ppt_nsteps=250, ppt_err_cut=0.08, with additional runs varying node counts and distance metrics (e.g., cosine). Pseudo-time was computed in scFates (85 parallel jobs, 100 mapping iterations, seed=42) and projected, together with the inferred tree, onto UMAP. Gene-trend clustering used a nearest-neighbor approach (n_neighbors=50, correlation metric, resolution=4), with outputs shown as heatmaps and single-gene trend plots.

### GSEA across temporal bins (trajectory-resolved enrichment)

To quantify pathway dynamics along trajectories, continuous pseudo-time t was binned at a width of 0.5 and intersected with Leiden clusters (Traag et al., 2019) to define robust subpopulations; bins with <50 cells were merged. Matrices were handled with Scanpy/Anndata (Wolf et al., 2018; Virshup et al., 2021). Ensembl → symbol mapping used MyGene.info (Xin et al., 2016).

For each (cluster × time) bin, we ran GSEApy (Fang et al., 2022) on MSigDB Hallmark 2020 with 100,000 permutations, minimum size = 10, maximum size = 100, and log₂ ratio of classes as the scoring metric. We report NES, *p*, FDR, and leading-edge genes, and summarize temporal trajectories of pathway activity across bins. Visual summaries (e.g., heatmaps/enrichment maps), as well as details of contrasts and multiple-testing handling, are included in the provided code.

### Survival analysis

Associations between gene expression and patient outcome were evaluated using the Kaplan–Meier Plotter (https://kmplot.com/analysis/), restricted to the pancreatic cancer dataset. Hazard ratios and log-rank p-values were retrieved for each gene shown in Fig. 4. The Kaplan–Meier Plotter integrates transcriptomic and survival data from multiple public repositories, enabling systematic validation of prognostic biomarkers (Gyorffy 2024a; Gyorffy 2024b).

### Network trajectory analysis

#### Data sources and preprocessing

Protein–protein interaction data were obtained from the OmniPath database (Türei et al., 2016) using the *omnipath* Python client v1.0.8. Only literature-curated interactions with directional and signed annotation were retained, ensuring a high-confidence starting network. For extracting a high-quality interaction, an omnipath curation effort of >=3 was chosen. All protein identifiers were standardized to UniProt IDs. Single-cell gene expression data were processed to identify expressed proteins in each experimental condition.

### Network reconstruction

#### Data source & global graph

Directed, signed protein–protein interactions were retrieved from OmniPath (omnipy client v1.0.8), keeping only literature-curated edges with curation effort ≥3 and standardizing IDs to UniProt (Türei et al., 2016). We intersected these interactions with the set of proteins expressed in our dataset (per condition) and assembled the global directed network in igraph v0.11.4, preserving the OmniPath type attribute (activation/inhibition/other); the largest connected component comprised N = 680 nodes and E = 1,403 edges (Csardi & Nepusz, 2006).

#### Group-specific subnetworks

For each experimental group (pseudo-time bin or treatment), we constructed an expression-filtered subnetwork by retaining only edges whose both endpoints were expressed (log(read count) > 0.5), then annotated nodes with gene symbols. Subnetworks spanned 119–259 nodes and 128–362 edges, with density 0.0053–0.0091 and average degree 2.15–2.80. An early bin (0_t_0.0000–0.5000) had 119 nodes / 128 edges (*density* = 0.009116; *avg_deg* = 2.151), whereas a later bin (2_t_2.5000–3.0000) had 259 / 362 (*density* = 0.005417; *avg_deg* = 2.795). Overall, node/edge counts increased with pseudo-time, with a slight density dilution but a higher average degree, indicating progressively strengthened pathway connectivity rather than mere network growth.

### Betweenness centrality across trajectories

For each trajectory-specific subnetwork (G1→S, TGF-β1, and the combined S-phase + TGF-β1 condition), we computed node betweenness centrality on the directed interaction graph, using shortest paths, to quantify how strongly each protein functions as a communication bottleneck. We then tracked betweenness values across the ordered experimental groups (pseudo-time) and summarized dynamics with simple slope-based trends (and corresponding Δ-changes), highlighting proteins whose centrality rose or fell substantially over the trajectory.

### Quantification of network-mediated perturbation influence

To assess how each protein contributes to the propagation of transcriptional perturbations across the interaction network, we quantified both structural and diffusive influence measures using Python (NetworkX, NumPy, Pandas). Protein–protein interaction networks were converted to undirected graphs, and pseudo-time-based perturbation bins were defined for *TGFB*, *CC*, and *Combined* treatment conditions.

For each bin, the 20 most perturbed genes (by absolute log₂ fold change, |log₂FC|) were designated as the *perturbed set*. Each network node was then evaluated for three complementary influence metrics:

1. Shortest-path betweenness fraction — the fraction of shortest paths between randomly sampled pairs of perturbed nodes that pass through the node. This reflects structural influence, identifying proteins that act as *bridges or bottlenecks* connecting active subnetworks.
2. Personalized PageRank (PPR) reach — computed by seeding PageRank with each node as the personalization source and summing steady-state probabilities over the perturbed set. This represents diffusive influence, i.e., the potential of a node to spread activity toward perturbed regions via all available paths.
3. Average shortest-path distance — the mean network distance from each node to all perturbed nodes, indicating proximity to transcriptionally active regions.

For each node and bin, these scores were integrated with |log₂FC| to calculate a composite influence score:

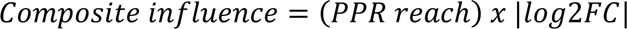

To detect synergy, scores in Combined were compared against TGFB + CC (sum): nodes with Composite_Combined > Composite_TGFB + Composite_CC, PPR_Combined > PPR_TGFB + PPR_CC, and log₂FC > 0 were retained. High-influence proteins were exported as tables (UniProt ID, gene symbol, composite score, fold-over-sum ratios, reach, betweenness, log₂FC).

### CDK1 path–overlap and randomized knockout analysis

We sought to identify upstream regulators that (i) participate in CDK1-mediated information flow and (ii) whose removal unloads the CDK1 bottleneck over pseudo-time. Analyses used directed protein–protein interaction (PPI) graphs built per trajectory bin (“trajectory graphs”) and one global directed PPI (“BIG graph”) used only for knockout (KO) neighborhood expansion. Computations were performed in Python using igraph (directed shortest paths and betweenness), NumPy, and Pandas. CDK1 was identified by UniProt P06493 and was never removed in any KO.

#### CDK1-mediated shortest-path overlap

Purpose: quantify whether a node v consistently co-occurs on shortest paths that pass through CDK1, i.e., whether v flanks or shares the CDK1 bridge.

Method: on the latest S-phase + TGF-β trajectory graph, we computed the all-pairs directed shortest-path distance matrix D in OUT mode. A pair (s, t) is CDK1-mediated if

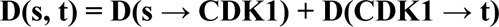

For each vertex v ≠ CDK1, we counted how many of these CDK1-mediated pairs also include v anywhere along the shortest path. We report overlap_count_latest (count) and overlap_frac_latest (fraction of all CDK1-mediated pairs). Graph size was below the exact all-pairs threshold (n < 3,000).

Interpretation: higher overlap_frac_latest indicates that v is a co-bottleneck or escort on CDK1-mediated routes.

#### Randomized knockout sweeps (multi-radius, multi-size)

Purpose: estimate the expected change in CDK1 betweenness when removing local neighborhoods around candidate regulators, and assess the robustness of effects.

KO universe comprised all nodes present in trajectory graphs (excluding CDK1). For each seed, a neighborhood on the BIG graph was expanded by OUT-neighbors to radius r ∈ {0, 1, 2}. KO set sizes were m ∈ {1, 2}. For each m, we ran 4,000 trials with a fixed random number generator (RNG) seed for reproducibility.

In each trial, the KO set was removed from every trajectory graph; directed betweenness for CDK1 was recomputed and averaged across graphs. The per-trial effect size was Δ = mean(B_CDK1^post) − mean(B_CDK1^baseline). Per-gene attribution assigned Δ/m to each seed in that trial; mean_delta is the average over all trials containing the seed. We also computed hit_rate_top1pct, the fraction of a seed’s trials in the top 1% by |Δ|.

Interpretation: mean_delta < 0 indicates that removing the candidate unloads the CDK1 bottleneck (reduces CDK1 betweenness); a high hit_rate_top1pct indicates a robust effect across random contexts.

#### Exact single-gene KO

Purpose: measure the deterministic effect of removing a single candidate on CDK1 betweenness.

Method: for each seed (typically r = 0), remove the seed from each trajectory graph and recompute directed betweenness; report singleKO_delta = mean(B_CDK1^post) − mean(B_CDK1^baseline).

Interpretation: singleKO_delta < 0 indicates that the seed alone unloads CDK1.

#### Aggregation and prioritization

Per-gene metrics were aggregated across radii r ∈ {0, 1, 2} (averaging effects; summing trial counts). We defined expected_effect = − mean_delta, so positive values indicate that a KO tends to reduce CDK1 betweenness.

A composite prioritization score was computed (overall KO score) as a weighted z-sum of four signals:

- overlap_frac_latest (0.35)
- expected_effect (0.35)
- hit_rate_top1pct (0.20)
- singleKO_delta (0.10)

Interpretation: Higher composite scores indicate higher-priority regulators that both sit on CDK1-mediated pathways and consistently unload the CDK1 bottleneck when perturbed. All computations used directed graphs; CDK1 was never removed.

### Simulated knockouts and CDK1 betweenness readout

To probe which upstream regulators sustain the CDK1/p21 bottleneck (Fig. 5A), we performed targeted knockouts (KOs) and recomputed directed betweenness centrality across the eight ordered trajectory-specific subnetworks (G1→S and S + TGF-β1), as described above. KO sets were built from regulators prioritized in Fig. 5B–C as seed ± ≤1 OUT-hop downstream neighbors on the global directed graph; each KO set was intersected with every trajectory subgraph and removed. CDK1 (P06493) was never removed. Effects were summarized as Δ betweenness (post-KO − baseline) for CDK1 per bin and by the slope over bins to capture temporal shifts.

### CDK1-centric network visualization

A CDK1-centered ego network was extracted from the global directed graph (k=2 upstream, k=1 downstream, ≤30 direct downstream by degree). The subgraph was rendered in NetworkX using a single union reference layout (Kamada–Kawai, seeded; CDK1 anchored at the origin) shared across panels. Panels overlay group-specific edges/nodes on a faint grey background; activating edges = green, inhibitory = red, other = grey dashed; arrows indicate direction.

### External validation in the PDAC single-cell atlas

#### Data source and preprocessing

To externally validate the bottleneck signatures identified in our 3D tissue model, we used the comprehensive PDAC single-cell atlas published by Loveless et al. (Clin Cancer Res, 2025), which integrates 726,107 cells from 231 patients across 12 independent studies (Zenodo accession 14199536). The atlas encompasses normal pancreatic tissue from healthy donors, adjacent-normal samples, treatment-naïve primary tumors, primary tumors resected after neoadjuvant chemotherapy (gemcitabine/abraxane, FOLFIRINOX, chemoradiation, RT + chemotherapy, or combined regimens), and metastatic lesions sampled from liver, peritoneum, and other sites. Cell-type annotations (ductal, cycling ductal, fibroblast, myeloid, lymphoid, endothelial, stellate, acinar, endocrine), molecular subtype classification (basal-like versus classical), cell-cycle phase assignments (G1, S, G2/M), and detailed treatment history were retained from the original atlas.

The atlas Seurat object (RDS format) was converted to AnnData (h5ad) format using SeuratDisk (v0.0.0.9020) in R (v4.3). The full-atlas UMAP coordinates were preserved for the multi-level visualization (Figure 6A–C). For epithelial-focused analyses, 300,577 cells annotated as ductal or cycling ductal populations were extracted and processed independently.

#### PHATE embedding

To preserve the continuous trajectory structure inherent in epithelial state transitions, we computed PHATE (Potential of Heat-diffusion for Affinity-based Transition Embedding; Moon et al., *Nature Biotechnology*, 2019) embeddings on the epithelial subset. Principal component analysis was first performed on the log-normalized expression matrix (50 components). PHATE was then computed using the phate Python package (v1.0.11) with default parameters (*k* = 5, *t* = ‘auto’, *n_components* = 2). The resulting two-dimensional embedding was used for all epithelial PHATE visualizations (Figure 6D–L, Figure 7).

#### Leiden signature scoring

Each epithelial cell was scored for the four Leiden-derived transcriptional signatures identified in our 3D PANC-1 tissue model: L0 (epithelial baseline), L1 (TGF-β/EMT program), L2 (S-phase + EMT bottleneck), and L3 (G2/M proliferative program). Signature scores were computed as the mean log-normalized expression of the top marker genes per Leiden cluster from the tissue model (19 genes for L2, variable for other clusters). The dominant Leiden identity for each cell was assigned based on the highest signature score.

To remove technical confounders, all continuous signature scores were corrected for library size by fitting a linear regression of the signature score on log(total UMI counts per cell) and using the residuals for downstream quantitative analyses, including AUC calculations and percentile-based enrichment. This correction ensures that differences in sequencing depth across studies and patients do not inflate signature scores.

#### CDK1–CDKN1A–WEE1 axis state classification

Binary expression states for *CDK1*, *CDKN1A*, and *WEE1* were defined using a >0 threshold on the raw count matrix (i.e., at least one detected UMI). Co-expression was defined as simultaneous detection of two or more axis genes in the same cell. Eight combinatorial states were constructed from all possible combinations of CDK1+/−, CDKN1A+/−, and WEE1+/− (denoted C+A+W+, C+A−W+, C+A+W−, C+A−W−, C−A+W+, C−A−W+, C−A+W−, C−A−W−).

#### Metastatic enrichment analysis (Figure 7A–C)

Metastatic association was quantified using Fisher’s exact test (two-sided) comparing the fraction of cells originating from metastatic lesions versus treatment-naïve primary tumors within each axis state or expression group. Odds ratios (OR) and associated p-values were computed using scipy.stats.fisher_exact (v1.11). The additive model (Figure 7A) tested successive addition of CDKN1A and WEE1 positivity among CDK1^+^ cycling cells (S + G2/M phase). The disease progression gradient (Figure 7C) computed CDK1^+^/CDKN1A^+^ co-expression rates separately for normal epithelium (donors + adjacent normal), primary tumor cells stratified by cell-cycle phase (G1, S, G2/M), and metastatic lesion cells.

#### Bridge mechanism analysis (Figure 7D–F)

To test whether CDKN1A co-expression functionally couples EMT and cell-cycle programs, we computed Spearman rank correlations between aggregate EMT and cell-cycle program scores across four mutually exclusive expression states defined by CDK1 and CDKN1A detection (Figure 7D). EMT scores were computed as mean expression of canonical EMT markers (*FN1*, *VIM*, *SERPINE1*, *TGFBI*, *SNAI2*, *MMP2*, *THBS1*); cell-cycle scores used S-phase and G2/M gene sets. Significance was assessed using scipy.stats.spearmanr with n > 4,000 cells per group.

Differential expression between CDK1^+^/CDKN1A^+^ co-expressing and CDK1^+^-only cycling cells (Figure 7E) was computed as log_2_ fold change of mean expression for selected EMT genes. Only cells in S or G2/M phase were included to control for cell-cycle confounding.

DUAL-program cells (Figure 7F) were defined as cells scoring above the 75th percentile for both the L1 (TGF-β/EMT) and L3 (G2/M) Leiden signatures simultaneously. DUAL-program membership rates were compared across CDK1/CDKN1A expression states and disease origins.

#### Signature specificity and AUC analysis (Figure 7G–I)

To compare the discriminative power of multiple gene signatures for metastatic versus treatment-naïve classification, receiver operating characteristic (ROC) analysis was performed using scikit-learn (v1.3.2). The following signatures were tested: (1) 27-gene bridge score (genes uniquely elevated in DUAL-program cells relative to both single-program populations), (2) 19-gene Leiden 2 bottleneck signature, (3) generic proliferation score (canonical S-phase + G2/M markers), (4) generic EMT score (Hallmark EMT gene set), (5) 50-gene bottleneck network score (from Figure 5 network analysis), (6) gemcitabine resistance gene panel (*GSTP1*, *ALDH1A1*, *ABCG2*, *ERCC1*, *RRM1*, *RRM2*), (7) general drug resistance panel, and (8) three-gene axis score (CDK1 + CDKN1A + WEE1). All scores were library-size corrected as described above prior to AUC computation. Area under the curve (AUC) was computed with 95% confidence intervals via bootstrapping (1,000 iterations).

Gene-level expression across conditions (Figure 7H) was visualized as a heatmap of log_2_ fold change relative to normal epithelium for selected axis and associated genes. Axis co-detection rates (Figure 7I) were computed as the fraction of cells co-expressing CDK1^+^/CDKN1A^+^ or all three axis genes (C^+^A^+^W^+^) within each condition, stratified by cell-cycle phase where indicated.

#### Treatment-specific analysis (Figure 7J–L)

Axis component prevalence among cycling cells (S + G2/M) was compared between treatment-naïve, gemcitabine/abraxane-treated, and FOLFIRINOX-treated groups (Figure 7J). Enrichment or depletion was quantified by Fisher’s exact test versus treatment-naïve cycling cells. Quiescence rates (Figure 7K) were computed as the fraction of cells in S + G2/M phase per treatment group; WEE1 detection among G1-arrested cells was overlaid as a secondary axis.

Percentile-based enrichment analysis (Figure 7L) tested whether treatment survivors and metastatic cells are enriched at the tail of the axis score distribution. For each percentile threshold (P50–P95 of the naïve primary cycling distribution), we computed the fraction of cells exceeding that threshold in gemcitabine cycling, FOLFIRINOX cycling, and metastatic groups, and expressed this as fold enrichment over the expected frequency in treatment-naïve cells.

#### Visualization and figure generation

All scatter plots (UMAP and PHATE) were generated using matplotlib (v3.8) with rasterized point rendering to maintain manageable file sizes despite >700,000 data points. Figure 6 (atlas mapping, 4 × 3 layout) and Figure 7 (quantitative validation, 4 × 3 layout) were exported at 1,200 DPI (PNG) and as vector PDF with fonttype 42 (editable text). Point sizes were scaled to cell density per panel to optimize visibility of rare populations (e.g., triple-positive cells). Yellow star markers highlight CDK1^+^/CDKN1A^+^ co-expressing cells in the treatment-specific panels (Figure 6J–L).

#### Statistical framework and limitations

All statistical tests were two-sided. Fisher’s exact tests were used for categorical comparisons (axis states versus disease origin) given the discrete nature of single-cell co-expression. Spearman correlations were used for continuous score comparisons. No multiple-testing correction was applied to the primary axis-state comparisons, as these represent hypothesis-driven tests of specific predictions from the tissue-culture model rather than exploratory genome-wide analyses.

We note that each treatment arm in the atlas derives from a limited number of studies and patients (gemcitabine/abraxane: predominantly one study; FOLFIRINOX: predominantly one study), and pre-treatment samples from the same patients are not available. Treatment-specific comparisons are therefore suggestive but cannot formally distinguish drug-induced selection effects from pre-existing patient-intrinsic or batch-related biology. Batch effects between contributing studies were not explicitly corrected, as we relied on the integrated embedding provided by the original atlas authors. All treatment-related interpretations are presented as observations consistent with, but not proof of, drug-specific selection.

#### Software and reproducibility

Analyses were performed in Python (v3.10) using scanpy (v1.9.6), phate (v1.0.11), scikit-learn (v1.3.2), scipy (v1.11), numpy (v1.24), and matplotlib (v3.8). Atlas conversion from RDS to h5ad used SeuratDisk (v0.0.0.9020) in R (v4.3). Code for all atlas validation analyses, figure generation, and statistical tests is available in the Supplementary Code repository.

## Results

### Systematic Single-Cell analysis of 3D PANC-1 tissue model: Cell cycle and TGF-β1-induced cell state cooperate in gemcitabine resistance

To investigate the mechanisms underlying Gemcitabine (GEM) resistance in a physiologically relevant context, we established 3D PANC-1 tissue models on decellularized porcine intestinal matrix (SISmuc), which recapitulate native tissue architecture and proliferation indices (Fig. 1A, D). Initial imaging revealed marked spatial heterogeneity, with subpopulations localized to surface regions and others embedded deep within crypt-like niches (Fig. 1 B). Eradication of PANC-1 cells by GEM could be assigned to crypt areas, suggesting microenvironment-specific differences in GEM sensitivity (Fig. 1B; HE staining). Quantification of Ki-67 immunofluorescent staining suggests that more proliferative and active surviving tumor cells remain after GEM treatment, particularly when stimulated by TGF-β1 (Fig. 1C). The MTT assay shows high chemoresistance against GEM in this 3D tissue model when examining the whole cell population (Fig. 1D).

**Figure 1.**
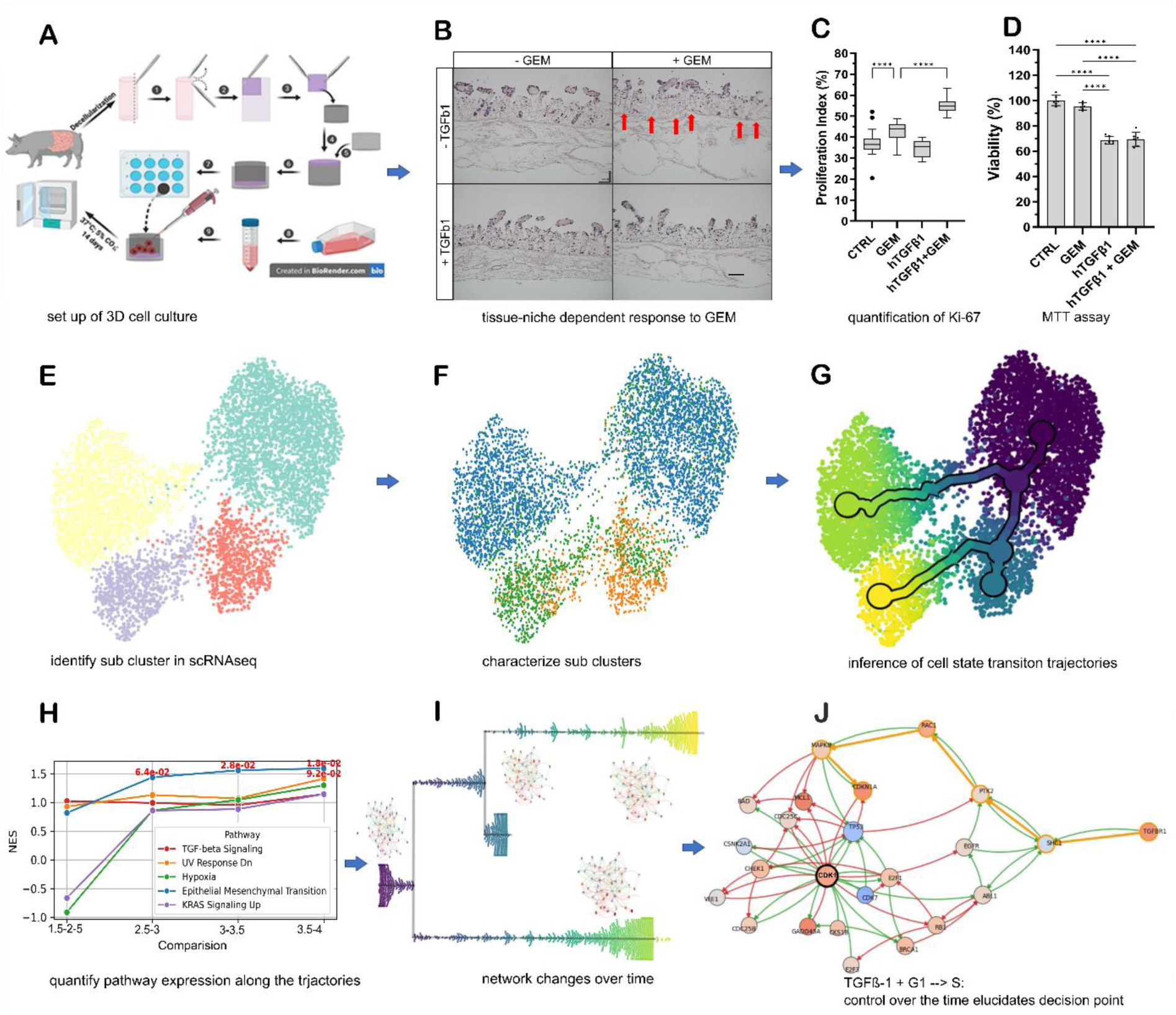
Integrated workflow for dissecting gemcitabine resistance in 3D PDAC cultures. (A) 3D PANC-1 cultures established on decellularized porcine intestinal matrix (SISmuc), recapitulating patient-like tissue architecture and proliferation indices. (B) H&E staining reveals spatial heterogeneity in GEM sensitivity (10 µM, 24 h): tumor cells in crypt niches are preferentially eradicated (red arrows), while TGF-β1-stimulated cells (11 days) invade across the preserved basement membrane. Scale bar: 100 µm. (C) Ki-67 quantification (5 images per condition) shows increased proliferation index after GEM treatment, further enhanced by TGF-β1 (n = 3; ****p < 0.0001). (D) MTT viability assay demonstrates high chemoresistance in the 3D model; TGF-β1 stimulation reduces metabolic activity, with no additional change upon GEM (n = 3; ****p < 0.0001, *p < 0.05). Error bars: SD. (E) Leiden clustering of scRNA-seq data identifies four subclusters. (F) Functional annotation assigns cell-cycle phase, EMT status, and pathway activities to each subcluster. (G) Pseudo-time trajectory inference reconstructs state transitions including G₁→S progression, EMT induction, and their intersections. (H) Gene and pathway dynamics along trajectories reveal state-specific drivers and co-regulated modules. (I) Gene regulatory networks reconstructed at baseline (G₁) and along each trajectory branch expose shifts in control architecture. (J) Centrality analysis and in silico knockout simulations identify the CDK1/CDKN1A axis as an emergent bottleneck unique to the combined G₁→S + TGF-β1–induced EMT state, pinpointing druggable upstream regulators.

Building on a companion bulk analysis that identified resistance-associated marker genes from this dataset (Caliskan et al., 2025), we applied single-cell RNA sequencing (scRNA-seq) to resolve the heterogeneity of the 3D cultures at high resolution. Unlike bulk analysis, which yields averaged expression profiles, scRNA-seq captures the molecular states of individual cells, revealing distinct subpopulations and gradual state transitions driven by cell-cycle progression, EMT, and their intersections (Figs. 1E-F).

Trajectory inference enabled the reconstruction of cell-state transitions within the 3D microenvironment, including developmental and stress-induced routes associated with resistance (e.g., G₁→S progression, EMT induction, partial G₂/M arrest) (Fig. 1G). By tracking gene and pathway expression dynamics along these trajectories, we identified state-specific drivers and co-regulated modules associated with particular PANC subclusters (Fig. 1H). In the following, we employ “dynamic” analyses to capture the evolving trajectories of gene expression over pseudo-time, in contrast to “static” approaches that assess expression states in isolation, such as through clustering or differential expression.

We then reconstructed gene regulatory networks for baseline quiescent (G₁) cells and for each major trajectory branch (G₁→S, S→G₂/M, S→EMT, G₁→EMT), enabling comparison of network control architecture across states (Fig. 1I). Centrality analyses revealed distinct shifts in control points, with the combined G₁→S + TGF-β1–induced EMT trajectory producing an emergent bottleneck centered on the CDK1/CDKN1A axis—absent in either program alone (Fig. 1J).

Upstream pathway mapping identified potential druggable regulators of these network control points, and in silico knockout simulations provided quantitative predictions for genes whose perturbation would most disrupt resistance-associated network topology. Together, this integrated spatial, transcriptional, and network framework provides a comprehensive mechanistic model of resistance in 3D PDAC cultures, generating experimentally testable hypotheses for targeted knockdown or other interference strategies (Fig. 1J).

### Cell cycle as driver of cell state heterogeneity: Static characterization of 3D PANC sub clusters

UMAP projections (Fig. 2A) clearly segregate the experimental conditions—CTRL, CTRL + GEM, CTRL + TGF-β1, and CTRL + GEM + TGF-β1—each forming two distinct clusters. We observe good quality control as replicates for CTRL (PANC, CTRL_1/2), CTRL + TGF-β1 (1+2), and CTRL + TGF-β1 + GEM (1+2) cluster together. Note that the CTRL + GEM condition lacks replicates due to the absence of data from cells that did not survive GEM treatment; therefore, this group should be interpreted with caution. To further delineate these subpopulations, we applied unsupervised Leiden clustering (Fig. 2B), which revealed distinct cell clusters and accounted for the high heterogeneity of the conditions, allowing us to investigate subclusters emerging in 3D tissue due to cellular differentiation. This analysis outlines pronounced differences in gene expression and pathway enrichment across clusters.

**Figure 2:**
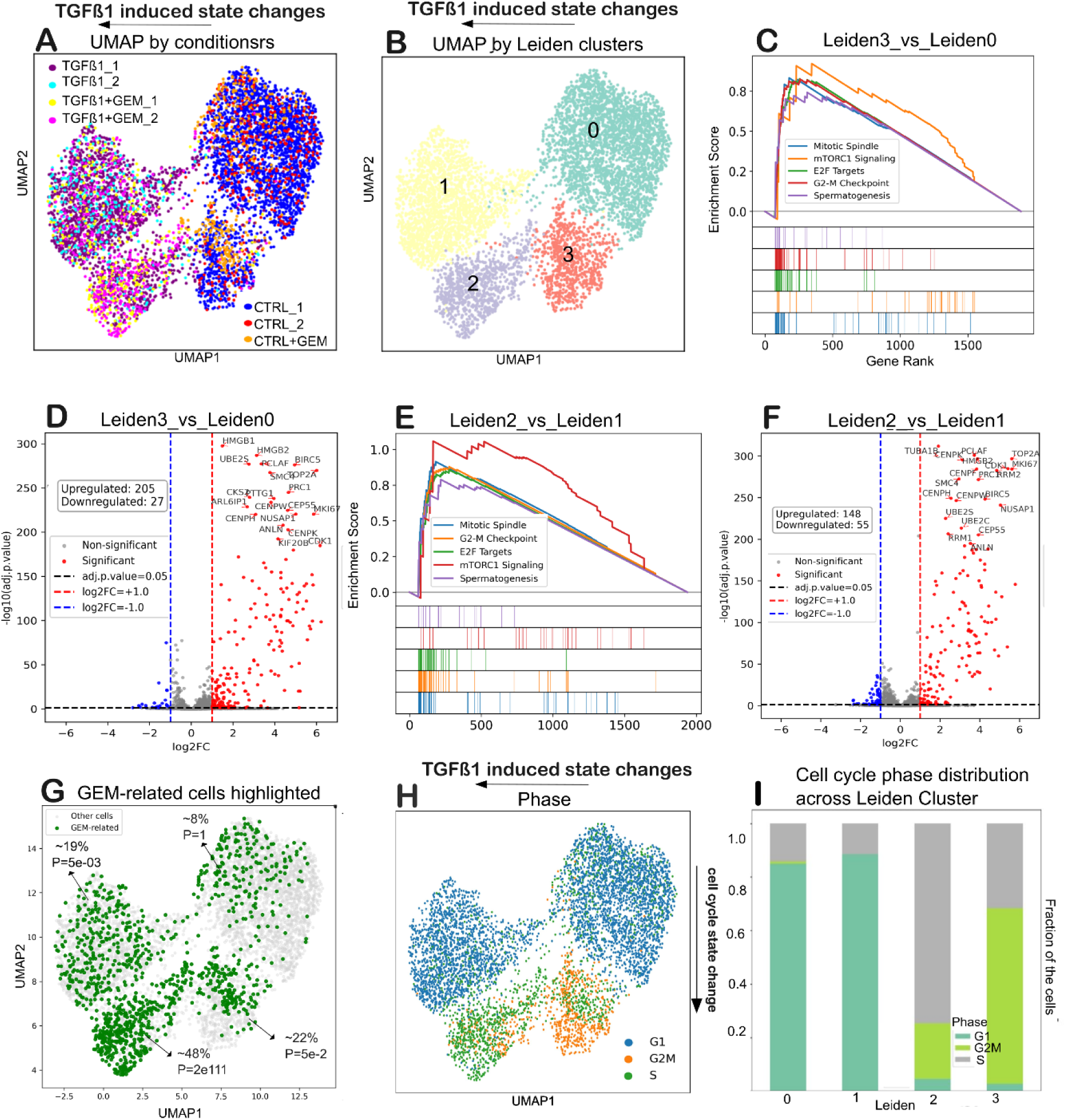
Subcluster Characterization of 3D PANC Tissues via Single-Cell Analysis. (A) UMAP projection of single-cell gene expression data for CTRL, CTRL + TGF-β, CTRL + GEM, and CTRL + GEM + TGF-β conditions. The projection reveals a heterogeneous expression landscape with cells organizing into distinct clusters. (B) Unsupervised Leiden clustering of the single-cell data delineates two major subclusters across the conditions, providing a refined categorization of the heterogeneous 3D tissue populations. (C) Gene Set Enrichment Analysis (GSEA) comparing Leiden cluster 3 versus cluster 0 (predominantly from CTRL and CTRL + GEM conditions) highlighting pathways related to transcriptional transition and cell cycle regulation (D) Volcano plots illustrate the differential gene expression between Leiden cluster 3 versus cluster 0, indicating the magnitude and direction of expression changes (E) GSEA comparing Leiden cluster 2 versus cluster 1 (from CTRL + TGF-β and CTRL + GEM + TGF-β conditions) (F) Volcano plots illustrate the differential gene expression between Leiden cluster 2versus cluster 1 (G) GEM-surviving cells highlighted in green across the UMAP. They distribute heterogeneously across the UMAP, while not forming distinct clusters. Enrichment analysis within Leiden clusters (with reported percentages and Fisher’s exact test p-values) identifies clusters with a higher proportion of GEM survivors. (H) Cell cycle phase distribution. UMAP embedding colored by predicted cell cycle phase (G1, S, G2M) showing the distribution of proliferative states across clusters in response to TGFβ1 ± GEM treatment. (I). Distribution of predicted cell cycle phases (G1, S, G2M) across Leiden clusters, revealing phase-specific enrichment in distinct clusters.

#### Heterogeneity of cell states within control (PANC/CTRL) cells

A comparison of Leiden cluster 3 versus cluster 0 (Fig. 2C) reveals significant enrichment in pathways related to mitotic spindle formation, E2F targets, and the G2-M checkpoint, indicating marked differences in cell cycle regulation. Differential expression analysis (Figs. 2D, S2A) identified approximately 200 upregulated and 30 downregulated genes between these clusters. Notable cell cycle regulators—including RRM2, CDK1, BIRC5, NUSAP1, CENPW, MKI67, CENPK, UBE2C, UBE2S, TK1, and HMGB1—were among the top differentially expressed genes, suggesting that shifts in cell division machinery underlie the static differences between these subclusters. (Note: Comparison of Leiden 2 vs. 3, Fig. S2B, reveals TGF-β1-induced changes with a very complex gene expression landscape (∼600 differentially expressed genes), warranting a more systematic evaluation.)

#### TGF-β-stimulation induces further heterogeneity of cell states

In cells stimulated with TGF-β1, subcluster comparisons (Leiden clusters 2 versus 1; Figs. 2E, 2F, S2C) reveal enrichment in similar cell cycle-related pathways, with approximately 148 genes upregulated and 55 downregulated between clusters. This static analysis reveals significant differences in pathways regulating mitotic progression and cell cycle checkpoints, which are crucial for the progression of cellular states under TGF-β1 stimulation. Although these data show distinct expression profiles, they predominantly highlight static pathway and gene expression differences affecting cell cycle regulation. (Note: Comparison of Leiden 1 vs. 0, Fig. S2D, reveals TGF-β1-induced changes with a very complex gene expression landscape (∼800 differentially expressed genes), indicating the need for further systematic evaluation.)

#### GEM treatment survivors do not form a distinct cluster

GEM treatment produces a more binary outcome, and GEM-surviving cells are integrated into existing Leiden clusters rather than forming a distinct one. Quantitative analysis of cell enrichment, assessed using Fisher’s exact test, indicates that surviving GEM-treated cells are distributed as follows: approximately 8% in Leiden cluster 0 (p ∼ 1), 20% in Leiden cluster 1 (p ∼ 5e-03), 48% in Leiden cluster 2 (p ∼ 2e-111), and 22% in Leiden cluster 3 (p ∼ 5e-02) (Fig. 2G). In Figures S3A-B, condition comparisons show that GEM-treated cells may share a common gene expression pattern involving genes such as H2AZ1, CENPW, TK1, RRM1, HELLS, CLSPN, RRM2, MYBL2, and NUSAP1, indicating roles in kinetochore regulation and cell cycle control. However, the high heterogeneity and numerous differentially expressed genes within the GEM-treated cell population (approximately 244 upregulated and 16 downregulated genes, as detailed in Figs. S3C and D) complicate the interpretation of the top changes. Many genes could serve as markers or contribute to tissue differentiation and GEM resistance.

#### Cell-cycle profiling reveals enrichment of proliferative states among GEM survivors

Cell cycle profiling validates transcriptional heterogeneity and highlights GEM-associated selection of proliferative states. Single-cell cell-cycle phase classification revealed strong enrichment of S and G2/M phase cells within Leiden clusters 2 and 3, whereas clusters 0 and 1 were predominantly assigned to G1 phase (Fig. 2H–I). Consistently, cell-cycle scoring and expression of key regulators such as CDK1 and AURKA (Fig. S4A–E) confirmed elevated cycling activity in clusters 2 and 3, in agreement with the transcriptional enrichment of E2F targets and G2/M checkpoint programs. These computational findings were supported by Ki-67 immunostaining, which demonstrated the persistence of proliferative cells following GEM exposure (Fig. 1D). Importantly, wet-lab quantification indicated a higher proportion of Ki-67-positive cells in GEM-treated samples compared to controls, suggesting that GEM preferentially eliminates sensitive cells while enriching for a resistant proliferative subpopulation. Notably, the combined TGF-β1 + GEM condition showed the strongest enrichment of Ki-67-positive survivors (∼55%), supporting the presence of a hybrid stress-tolerant and proliferative phenotype.

Overall, our static analysis of 3D cell populations under control, GEM, and TGF-β1 conditions highlights pronounced differences in pathways governing cell cycle progression and mitotic regulation. The significant heterogeneity observed within and across Leiden clusters, as well as the substantial expression differences in GEM-treated populations, underscores the need for a more granular analysis of expression dynamics to elucidate the regulatory mechanisms underlying tissue differentiation and chemoresistance fully.

Transitioning from static Leiden clustering to dynamic cell state analysis, we leverage the inherent power of single-cell data to reconstruct continuous cell state trajectories and infer pseudo-time. Unlike bulk comparisons that yield averaged signals, single-cell analysis captures the full diversity of individual cellular states, enabling us to sort cells along a continuum that reflects progressive gene expression changes. For instance, by ordering cells from an early state (observed in Leiden cluster 0) to a more advanced or treatment-associated state (as seen in Leiden cluster 2, which is exclusive to the 3D tissue context), we delineate the gradual transcriptional changes underlying tissue differentiation. This approach is particularly pertinent for both control PANC cells and TGF-β1-induced populations, as TGF-β1 stimulation triggers a cascade of gene expression alterations—from early direct targets to later differentiation markers—implicated in aggressive cancer phenotypes. By integrating inference with differential gene expression (as presented in Figs S2 and S3), we validate key genes driving these transitions and further substantiate our findings through survival analyses of candidate targets. In this manner, dynamic trajectory reconstruction not only complements static clustering but also deepens our understanding of the regulatory mechanisms underlying TGFβ-1 induction and chemoresistance (Figure 3).

**Figure 3:**
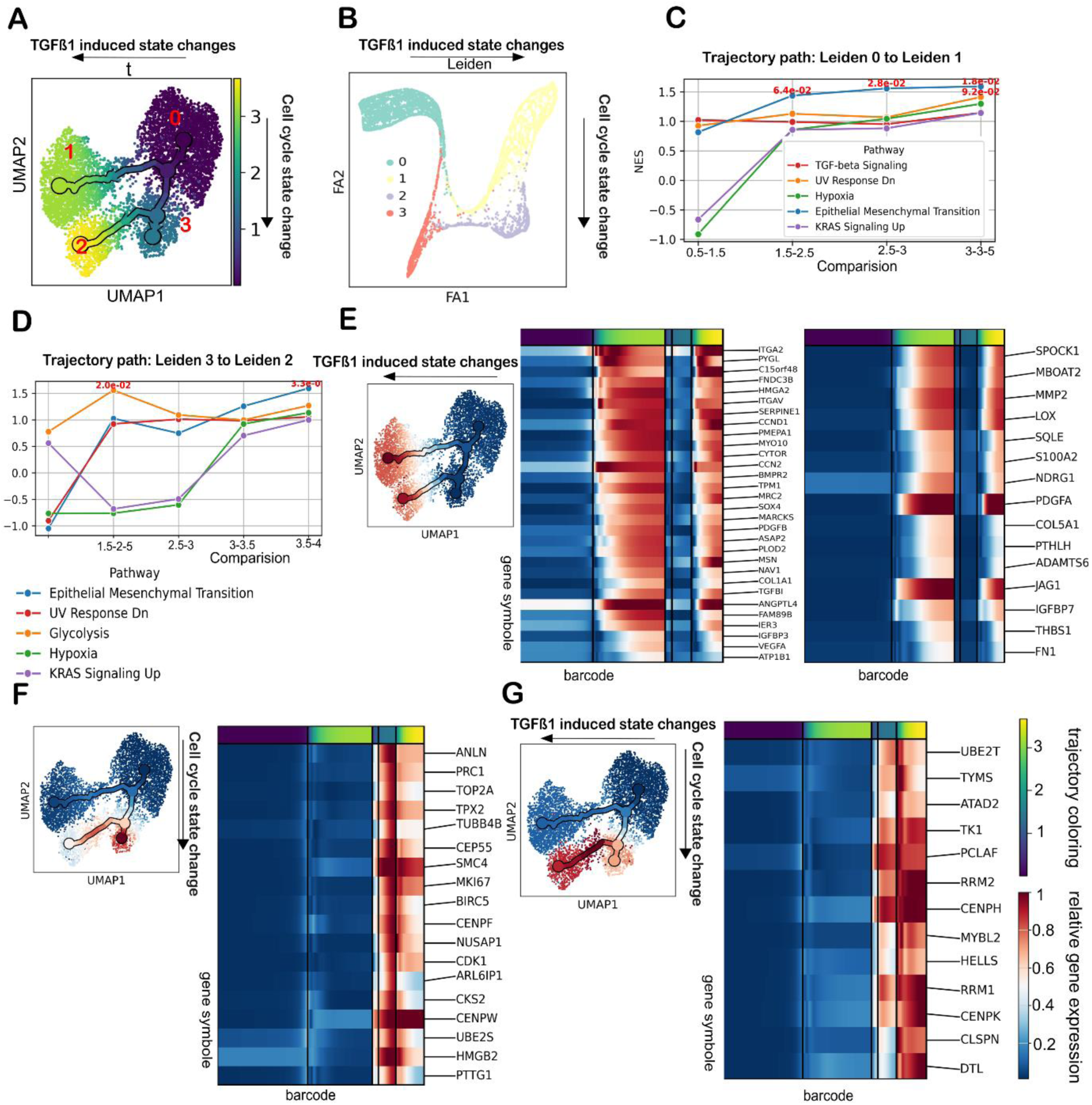
Profiles of state change trajectories in 3D PANC tissues. (A) UMAP visualization of single-cell transcriptomes from 3D PANC tissues, with cells ordered by pseudo-time (violet: early [0]; yellow: late [3.7]), red numbers show Leiden clusters. Two distinct trajectories are observed, originating from control subpopulations and progressing to late states upon TGF-β1 treatment. (B) Force atlas representation of the same dataset, with cells color-coded by Leiden cluster, illustrating continuous transitions and bridging cell states along the differentiation trajectories. (C-D) Temporal pathway profiling was performed using Gene Set Enrichment Analysis (GSEA) with the Hallmark gene set. Two trajectories were analyzed: one from the major PANC-1 sub-group (Leiden cluster 0) to the bigger TGF-β stimulated sub-group (Leiden cluster 1) and from the smaller control sub-group (Leiden cluster 3) to the smaller TGF-β1–treated sub-group (Leiden cluster 2). Normalized enrichment scores (NES) are presented across binned pseudo-time, showing a stepwise activation of key signaling pathways associated with TGF-β1 stimulation. (E–G) Sequential gene expression profiling was used to cluster dynamic expression patterns along the trajectories into three distinct groups: (E) one group representing genes sequentially induced by TGF-β1; (F) a second group reflecting changes in cell cycle state; (G) and a third group showing overlapping expression patterns from both processes. UMAP mean expression profiles serve as a baseline control for trajectory inference. Color gradients in heatmaps represent relative gene expression (scaled 0–1) along pseudo-time, while trajectory color bars denote pseudo-time progression (0–3.75). Corresponding downregulated gene sets are provided in the Supplementary Figures S2 and S3.

### Dynamic reconstruction of cell state transitions in 3D tissue

In Figure 3, we extend our static Leiden clustering analysis (Figure 2) by reconstructing continuous cell state trajectories. Single-cell analysis enables us to order individual cells along a continuum based on their gene expression profiles, thereby capturing the gradual transitions in cellular states. Cells are arranged such that each cell is closely related to its immediate neighbor with a subtly shifted transcriptional profile, allowing us to infer stepwise changes that reflect the progression from a source to a target state.

For control PANC-1 cells, one distinct root populations are evident, corresponding to Leiden cluster 0, characterized by low expression of E2F targets, mitotic spindle components, and associated cell cycle pathways (see ForceAtlas representation in Fig. S5A). This root serves as the starting point for major transcriptional trajectories. As cells transition from this root, they acquire gene expression profiles characteristic of more advanced states (Fig. S5B). From the two CTRL subclusters to the two TGF-β1 subclusters, two similar trajectory patterns are observed that develop in parallel, with gradual shifts in gene expression reflecting the sequential activation of early TGF-β1 targets and later transcriptional markers (Fig. S5D). The inferred pseudo-time metric, though relative and not an absolute time measure, reveals regions of steep transcriptional change as well as more gradual transitions along these trajectories.

Figure 3A displays the primary trajectories overlaid on a UMAP projection, with pseudo-time values color-coded to indicate progression from early states (Leiden 0 and 3) to more differentiated states (Leiden 1 and 2). Figure 3B presents a force atlas visualization, emphasizing the continuity of the expression landscape and the bridging cell populations that connect the discrete clusters. The Leiden clusters are colored, indicating a good overlap between the Leiden classification and the inferred temporal trajectories.

Figure 3C illustrates the dynamic expression of pathways over pseudo-time, as cells progress from CTRL (Leiden cluster 0) to CTRL + TGF-β1 (Leiden cluster 1), representing the putative less GEM-resistant cell population. The normalized enrichment scores (NES) demonstrate that TGF-β1 stimulation induces a persistently high level of TGF-β1 signaling, accompanied by a significant activation of epithelial-to-mesenchymal transition (EMT) and UV response pathways—both hallmark indicators of TGF-β1 activity. Additionally, pathways related to hypoxia and Kras signaling are upregulated, although their statistical significance remains less definitive. In Figure 3D, a similar trend is observed along the second trajectory, confirming the robust upregulation of EMT, glycolysis, and hypoxia pathways, with Kras signaling also elevated, albeit without reaching clear significance thresholds.

Our dynamic clustering of gene expression profiles along the inferred trajectories reveals three major groups that elucidate the regulatory programs driving cell state transitions. The overall UMAP expression of these gene clusters serves as a baseline control for our trajectory inference. In addition, as demonstrated in Figure 2, the differential expression patterns between PANC (CTRL) and TGF-β1 subpopulations are recapitulated in these dynamic analyses, providing a strong validation of our static and dynamic approaches.

Figure 3E highlights a set of genes sequentially induced by TGF-β1, organized according to their temporal activation during differentiation. Early response genes—including TGF-β1, SERPINE1, MYO10, TPM1, HMGA2, PDGFB, C15orf48, IER3, VEGFA, and ATP1B1—are activated at initial stages. As cells progress along the trajectory, a later wave of expression is observed, with markers such as COL1A1, PDGFA, MBOATT2, LOX, and SPOCK1 emerging, followed by a subset with very late onset (PGM2L1, ADAMTS6, COL5A1, S100A2, PTHLH, NDGR1, THBS1, and FN1). This ordered induction narrows down hundreds of genes to a focused group of candidates that are critical for driving specific cell state transitions and the TGF-β1–mediated differentiation process.

Figure S6 confirms that genes such as KRT8, KRT18, KRT19, HMGB3, UBE2C, CAV1, and CD55 are progressively downregulated during TGF-β1–mediated differentiation, reflecting a coordinated loss of epithelial identity. This observation is consistent with our static differential expression analysis (Figure 2), further validating our findings.

In Figure 3G, we identify an intriguing cluster of gene expression profiles along the trajectories that exhibit overlapping patterns indicative of both cell cycle changes and TGF-β1–induced state transitions. These genes, which are highly expressed in GEM–survivor–enriched populations (notably in Leiden clusters 2 and 3), may serve as key markers—or even drivers—of a combined chemoresistance phenotype. Interestingly, the overall UMAP expression pattern of these genes mirrors that observed in Figure 2G (enrichment of GEM survivors). This suggests a potential connection between GEM survival and the gene expression program seen in Figure 3G. Specifically, GEM-treated cells appear enriched across transcriptional states associated with cell cycle changes, TGF-β1–induced state transitions, and a combination of both. GEM enrichment percentages further support this link: approximately 22% of GEM-treated cells localize to the cell cycle–related cluster (Leiden cluster 3), ∼19% to the TGF-β1–induced cluster (Leiden cluster 1), and the highest proportion (∼48%) to the combined state represented by Leiden cluster 2.

The detailed analysis is given in Supplementary Material, Figure S1-S7 together with extended methods and further discussion. In particular, some of the top genes that appear to be common in all GEM survivors (CTRL + GEM vs. CTRL), as shown in Figure S3A, are present in this cluster. These findings imply that GEM resistance may be influenced by the interplay of these transcriptional programs rather than arising from a distinct, newly formed gene expression state.

Together, these results (Panels E–G) provide a comprehensive view of the sequential gene expression changes driving cell state transitions. They offer mechanistic insights into the processes of cell cycle and TGF-β1–mediated transition, and suggest that the overlapping gene signatures may also be involved in the development of chemoresistance. Finally, Table S2 presents a brief Principal Feature Analysis (PFA) that further supports these findings by quantifying the information content of key regulatory genes, reinforcing their critical roles in the observed differentiation cascades.

### Key marker genes of defined cell states linked to poor clinical prognosis

Dynamic trajectory analysis of single-cell data resolved three distinct gene groups driving transcriptional changes in 3D PANC-1 tissue models (Fig. 4). Group A genes were sequentially induced during the TGF-β1–driven transition, Group B genes reflected cell cycle–associated changes during G1→S progression, and Group C genes exhibited overlapping expression patterns from both TGF-β1 and G1→S signatures. For each group, representative genes are shown with two complementary visualizations: (i) UMAP projections with expression levels color-coded (left panels) and (ii) inferred expression trajectories fitted along pseudo-time (right panels). This side-by-side layout ensured consistency between computational inference and observed cellular expression patterns. To assess clinical relevance, we integrated survival data from the Kaplan–Meier Plotter database (https://kmplot.com/analysis/) focusing on pancreatic cancer. Hazard ratios are shown within the figure panels, demonstrating that several genes across all three groups are significantly associated with poor outcomes of patient survival.

**Figure 4.**
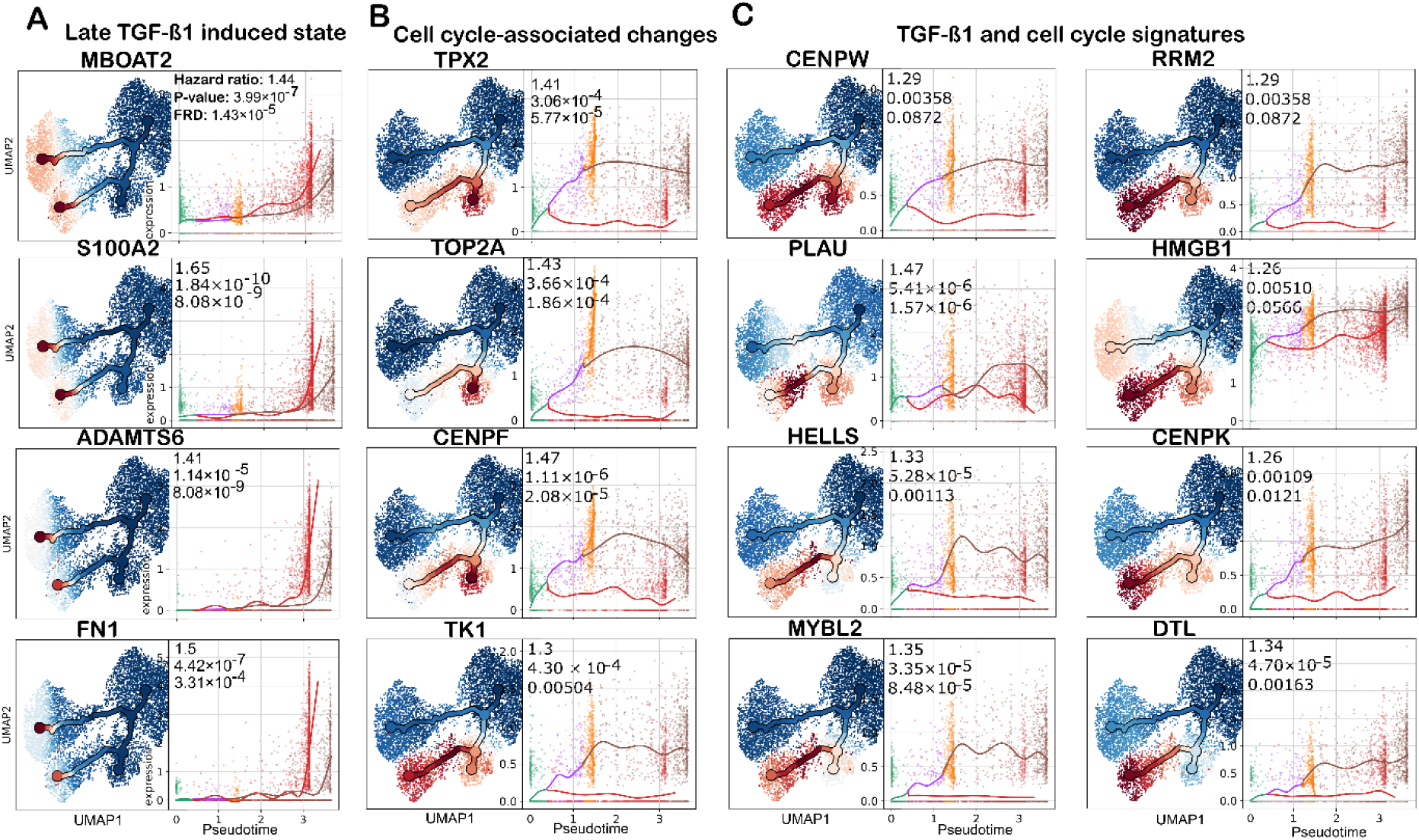
Key marker genes defining distinct cell states associated with poor clinical prognosis. (A–C) UMAP and pseudo-time plots showing the expression patterns of selected genes representing distinct transcriptional modules in 3D-cultured PANC-1 cells following TGF-β1 stimulation. Each gene panel includes a UMAP embedding (left) with cells colored by fitted expression values (red = high, blue = low) and the corresponding pseudo-time trend plot (right) showing gene expression dynamics along the inferred trajectory branches. Each color in the pseudo-time plots corresponds to a milestone along the Leiden-defined clusters: green – early milestone (Leiden 0), orange – intermediate milestone (Leiden 3), red – late milestone (Leiden 1), and brown – late milestone (Leiden 2). Solid colored lines represent the average smoothed expression trends for each branch. (A) Representative marker genes exhibiting late-onset activation following TGF-β1–induced state transitions. (B) Genes associated with cell cycle–driven changes, marking G1→S phase progression. (C) Genes showing combined expression signatures of TGF-β1 induction and cell cycle regulation, reflecting overlapping transcriptional programs that sustain these transition processes. Hazard ratios (HR), P-values, and FDR-adjusted significance levels from Kaplan–Meier survival analyses are provided to indicate associations with clinical outcomes.

#### Group A – Sequentially induced by TGF-β1 signaling

This group comprises genes that are progressively activated during TGF-β1–mediated signaling, typically with a late onset along the pseudo-time trajectory. Examples include *MBOAT2*, *S100A2*, *PGM2L1*, and *FN1*. Survival analysis yielded HRs of approximately 1.44, 1.65, 1.41, and 1.50, respectively, highlighting their association with poor prognosis (Fig. 4A).

#### Group B – Cell cycle–associated changes (G1→S transition)

Genes in this group mark proliferative heterogeneity within the 3D PANC-1 model, reflecting G1→S progression. Key examples include *TPX2*, *TOP2A*, *CENPF*, and *TK1*, with HRs of approximately 1.41, 1.43, 1.47, and 1.30 (Fig. 4B). These genes have been previously linked to gemcitabine (GEM) resistance and cell cycle regulation.

#### Group C – Overlapping TGF-β1 and G1→S signatures

Genes in this category combine features of both TGF-β1 induction and cell cycle–associated expression and are enriched in GEM-survivor populations. Examples include *HELLS*, *MYBL2*, *PLAU*, and *DTL*, with HRs of approximately 1.33, 1.40, 1.35, and 1.47 (Fig. 4C). Their consistent association with adverse outcomes suggests roles in chemoresistance and tumor persistence.

### A coupled S-phase + TGF-β1 state concentrates network control in CDK1/p21 and reveals licensing/checkpoint controllers

We asked how network control shifts along three trajectories—G1→S (cell-cycle), TGF-β1 (EMT-like), and the combined S-phase + TGF-β1 state linked to gemcitabine survival.

The analysis of betweenness centrality in Figure 5A illustrates how network nodes serve as bridges between distinct modules across various trajectories. The G₁→S cell cycle transition trajectory demonstrated increased betweenness centrality for cell–cycle–related nodes, including CDK1, EGFR, and PRKCA. The TGF-β1-induced EMT trajectory showed a slight increase in betweenness centrality for RHOA, ROCK1, EZR, PRKCA, and additional cytoskeleton regulatory nodes. The combined S-phase + TGF-β1 condition (cluster 2) produced an exceptional increase in betweenness centrality for CDK1 and CDKN1A, which exceeded the values of both individual trajectories (Δbetweenness ≈ 5000 for CDK1 and ≈ 4000 for CDKN1A; slopes in Fig. 5A). The sudden importance of CDK1 and CDKN1A emerges as critical connectors that unite different modules when S-phase progression and TGF-β1-induced EMT occur together. The large number of genes affected by TGF-β1 and S-phase induction (Figures 2 and 3) leads to this centrality peak because CDK1 and CDKN1A connect two extensive independent modules that operate through cell-cycle programs and EMT/TGF-β1 signaling pathways.

**Figure 5.**
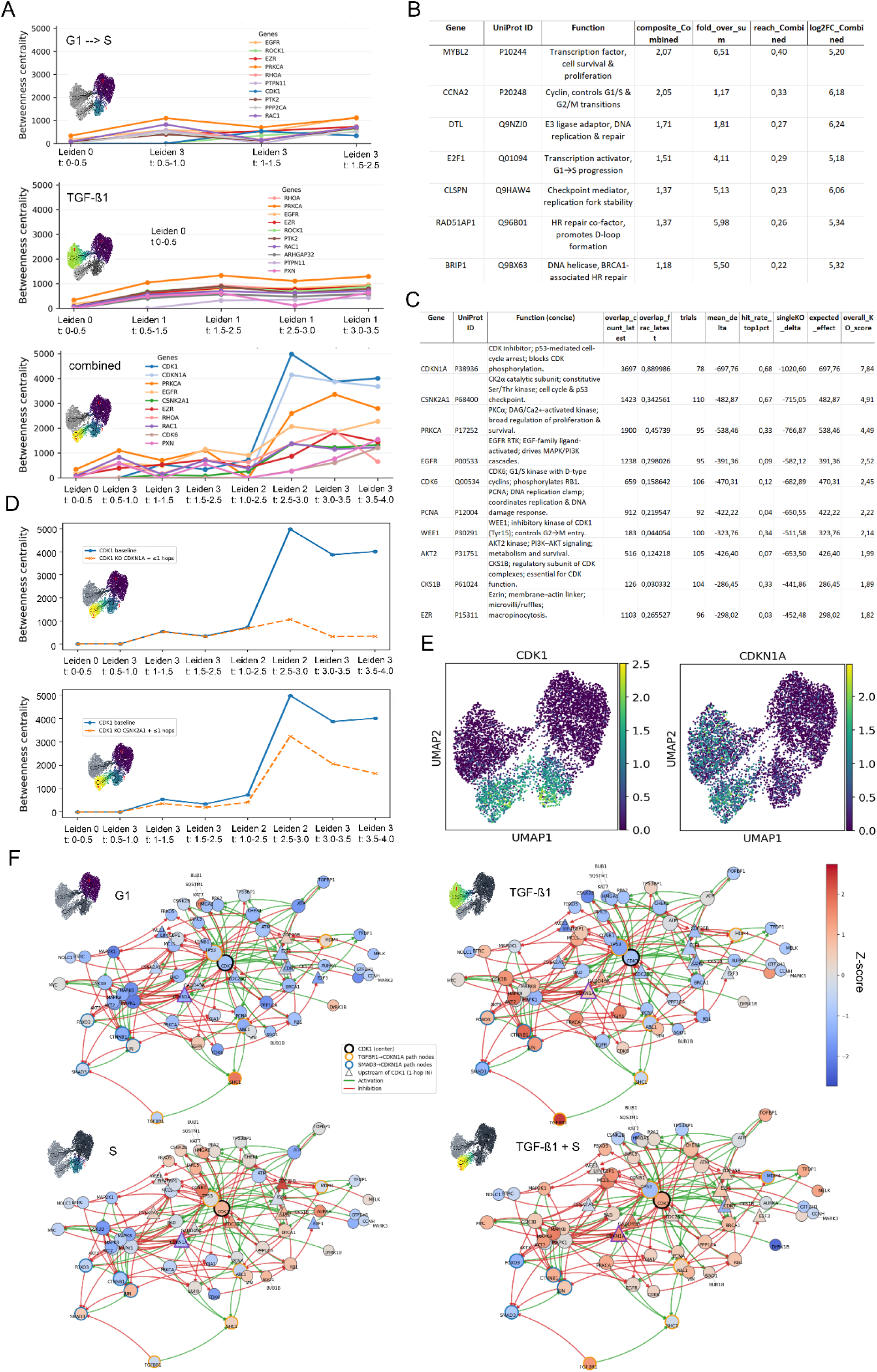
Network determinants of the Combined (G1→S + TGF-β1) state. (A) Betweenness profiles of network nodes along the G1→S, TGF-β1, and Combined trajectories across pseudo-time. (B) Genes with high network-mediated influence in the Combined state. Columns: Gene symbol, UniProt ID, Function, composite_Combined, fold_over_sum, reach_Combined, log2FC_Combined. The composite score is (PPR reach) × |log₂FC| computed on the undirected PPI (α=0.85; top-perturbed N=20). See Methods for full details. fold_over_sum = composite_Combined / (composite_TGFB + composite_CC) summarizes synergy: values >1 highlight genes that are unusually influential in the Combined state compared with G1→S or TGF-β1 alone. We display only entries passing the synergy filter (composite_Combined > composite_TGFB + composite_CC, reach_Combined > reach_TGFB + reach_CC, log2FC_Combined > 0). (C) CDK1 co-bottlenecks and KO sensitivity. Table ranks upstream nodes by (i) their co-occurrence on CDK1-mediated shortest paths in the latest Combined bin (overlap_count_latest, overlap_frac_latest) and (ii) how much their removal unloads the CDK1 bottleneck across trajectories (trials, mean ΔCDK1 betweenness, hit_rate_top1pct, singleKO_delta). Higher overlap_frac_latest marks frequent “escorts” on s→CDK1→t routes; negative mean Δ or singleKO_delta indicates unloading. (If shown) overall_KO_score integrates these signals (weights: 0.35 overlap_frac_latest, 0.35 expected_effect=−mean_delta, 0.20 hit_rate_top1pct, 0.10 singleKO_delta). (D) Betweenness of CDK1 across pseudo-time (G1→S and TGF-β1 trajectories) before/after knockout of CDKN1A and CSNK2A1 and its direct downstream interactors. (E) Reference expression of CDK1 and CDKN1A is shown in a UMAP. (F) CDK1-anchored network views across four states (G1, S, TGF-β1, S+TGF-β1). Edges are encoded by color/style: green = activating, red = inhibitory, grey dashed = other; arrows indicate direction. Node color encodes the expression z-score of the log expression in that state (diverging scale, globally normalized across panels). CDK1 is emphasized (thicker border/outline). Labels show gene symbols.

#### A coupled S-phase + TGF-β1 state concentrates network control in CDK1/p21

We followed how betweenness centrality shifts along three paths—G1→S (cell-cycle), TGF-β1 (EMT-like), and their combined state linked to gemcitabine survival. In G1→S, classic cell-cycle players (CDK1, EGFR, PRKCA) move to the center. Under TGF-β1 alone, EMT/cytoskeletal nodes (RHOA, ROCK1, EZR, PRKCA) rise modestly. Strikingly, in the combined condition, we see a sharp spike specifically for CDK1 and CDKN1A/p21 (Δbetweenness ≈ 5000 and 4000, respectively; slopes in Fig. 5A). In other words, only when S-phase progression and TGF-β/EMT run together do CDK1 and p21 become the bridge that ties the two modules. Many shortest paths now funnel through CDK1/p21, creating a bottleneck that correlates with the enrichment of survivors treated with Gemcitabine.

The CDK1/p21 bottleneck in Fig. 5A indicates that, when S-phase and TGF-β1 co-occur, control is funneled through a narrow bridge linking cell-cycle and EMT/TGF-β modules. We next asked which genes, in this joint state, exert unusually broad influence over the differentially expressed set. Using a simple diffusion-plus-amplitude score (composite influence = PPR reach × |log₂FC|) and a synergy filter (fold_over_sum > 1; see Methods), we ranked genes in the Combined condition.

#### From rewiring to influence: genes amplifying the CDK1/p21 bottleneck

Consistent with the CDK1/p21 isthmus in Fig. 5A, the high-influence set in the combined S-phase + TGF-β1 state resolves into three modules that converge on that gate from opposite sides:

- G1/S transcriptional push (cell-cycle arm): MYBL2 (P10244) and E2F1 (Q01094) show strong synergy (fold_over_sum ≈ 6.5 and ≈ 4.1) and high reach (≈ 0.40–0.29), driving expression of S-phase/mitotic targets; CCNA2 (P20248) (composite ≈ 2.05; modest synergy ≈ 1.17) activates CDK1 complexes, amplifying the pro-mitotic flow toward the gate.
- Licensing/checkpoint brake (checkpoint/TGF-β arm): DTL (Q9NZJ0) (CRL4^CDT2; synergy ≈ 1.81) and CLSPN (Q9HAW4) (ATR→CHK1; synergy ≈ 5.1) load the gate from the restraint side—DTL modulating the p21/CDT1 axis during S-phase, CLSPN feeding CHK1-mediated CDC25 inhibition that suppresses CDK1 activation.
- Damage-tolerance support (stabilizers): RAD51AP1 (Q96B01) and BRIP1 (Q9BX63) (synergy ≈ 6.0 and ≈ 5.5; reach ≈ 0.26–0.22) provide homologous-recombination(HR)/Fanconi support(FA), helping the network sustain S-phase pressure on the gate under stress.

Operationally, reach captures how broadly a gene can diffuse influence toward many perturbed targets in the Combined state, while fold_over_sum > 1 marks influence that is amplified specifically under co-occurring S-phase and TGF-β1 rather than simply additive. Together with Fig. 5A, these results indicate that a pro-mitotic push (E2F1/MYBL2/CCNA2) is counter-weighted by a checkpoint/licensing brake (CLSPN→CHK1, DTL/p21), with HR/FA nodes stabilizing the state—all converging on the CDK1/p21 gate in the S-phase + TGF-β1 condition.

Building on the state-specific CDK1/p21 bottleneck seen in the S-phase + TGF-β1 condition, we next asked which upstream regulators both sit on CDK1-mediated routes and actually collapse CDK1’s bridging role when removed. In Fig. 5C, two candidates clearly dominate: CDKN1A/p21 and CSNK2A1/CK2α. For p21, the overlap count is ∼3,697 with an overlap fraction of ∼0.890—meaning roughly nine out of ten shortest paths that traverse CDK1 also pass through p21 in the latest Combined bin. When p21 is knocked out, CDK1’s betweenness drops sharply (mean Δ ≈ −698 across 78 trials; exact single-gene KO Δ ≈ −1,021), and this effect is robust (top-1% hit-rate ≈ 0.68), yielding the top overall_KO_score (∼7.84). CK2α shows a similar, if smaller, profile: overlap count ∼1,423, overlap fraction ∼0.343, and KO-driven unloading of CDK1 (mean Δ ≈ −483 across 110 trials; single-KO Δ ≈ −715; hit-rate ≈ 0.67; overall_KO_score ∼4.91). In plain terms, the overlap metrics capture how often a gene co-occurs on CDK1-mediated shortest paths (co-bottleneck strength), while mean_delta and singleKO_delta quantify how much CDK1’s centrality falls when that gene is removed (negative values indicate unloading). The hit_rate_top1pct reflects the stability of this effect across randomized contexts, and the overall_KO_score summarizes these signals into a single priority readout.

Figures 5D–E put these numbers into biological context. Removing CDKN1A or CSNK2A1 consistently lowers CDK1 betweenness across pseudo-time, demonstrating that they sustain the gateway. Expression patterns align with this mechanism: CDK1 rises along the G1→S trajectory, whereas CDKN1A is induced by TGF-β1 rather than by S-phase. Their co-occurrence—S-phase pressure increasing CDK1 abundance and a TGF-β1-driven brake via p21—creates the narrow isthmus where the two programs meet, explaining the sharp betweenness spike and identifying p21 and CK2α as the most proximate drivers of the CDK1/p21 control point.

The network visualization shows that CDK1 acts as a connection point unifying the cell-cycle/mitotic module with the TGF-β1/SMAD signaling module (Fig. 5F). The cell-cycle module operates during the S-phase but exhibits minimal interaction with EMT signaling pathways. The EMT module shows activity in the TGF–β1–only condition but lacks strong connections to cell cycle regulation. The combined S-phase + TGF-β1 state enables CDK1 to create multiple direct connections between these major modules, which results in its increased betweenness centrality and essential function in the resistant phenotype. The network nodes that show both significant expression changes and extensive network connections (DTL, MYBL2, CLSPN, RAD51AP1, and CCNA1) maintain close relationships with CDK1 hubs and their downstream elements, thus supporting CDK1’s function as a coordinator between cell proliferation and EMT-related processes.

### The CDK1–CDKN1A–WEE1 bottleneck signature maps onto a comprehensive PDAC single-cell atlas

Our trajectory-based network analysis in TGF-β1-treated pancreatic ductal cells identified a regulatory bottleneck connecting two otherwise antagonistic transcriptional modules: an S-phase/cell-cycle program driven by *CDK1*, *RRM2*, and *TOP2A*, and a TGF-β1-induced EMT program anchored by *FN1*, *SERPINE1*, and *TGFBI*. The bottleneck axis—defined by the simultaneous expression of *CDK1* (cell-cycle driver), *CDKN1A* (paradoxical bridge), and *WEE1* (G2/M checkpoint brake)—was identified in a controlled tissue-culture model using dynamic single-cell trajectory analysis. To determine whether this bottleneck represents a biologically meaningful state in human disease, we sought to validate these signatures in an independent, large-scale clinical resource.

We leveraged a recently published comprehensive pancreatic ductal adenocarcinoma (PDAC) single-cell atlas integrating 726,107 cells from 231 patients across 12 independent studies (Loveless et al., 2025). This atlas encompasses the full spectrum of PDAC disease states, including normal pancreatic tissue from healthy donors and adjacent-normal samples, treatment-naïve primary tumors, primary tumors resected after neoadjuvant chemotherapy (gemcitabine/abraxane, FOLFIRINOX, chemoradiation, or combined regimens), and metastatic lesions sampled from liver, peritoneum, and other sites. The atlas provides a rich annotation framework including cell-type classification (ductal, fibroblast, myeloid, lymphoid, endothelial, stellate, acinar, and endocrine populations), molecular subtype assignment (basal-like versus classical), cell-cycle phase scoring (G1, S, G2/M), and detailed treatment history.

From the full atlas, we extracted 300,577 epithelial cells (ductal and cycling ductal populations), which constitute the relevant compartment for PDAC tumor biology. We computed PHATE embeddings to preserve the continuous trajectory structure inherent in epithelial state transitions, and scored each cell for the four Leiden-derived signatures from our tissue-culture model: L0 (epithelial baseline), L1 (TGF-β/EMT program), L2 (S-phase + EMT bottleneck), and L3 (G2/M proliferative program).

#### Multi-level mapping reveals bottleneck localization in the atlas

To orient these signatures within the broader tissue context, we performed a multi-level analysis. At the full-atlas level (Figure 6A–C), Leiden signature scoring of the epithelial compartment revealed that L2 (bottleneck) cells occupy a spatially defined region on the UMAP embedding, concentrated at the interface between ductal and cycling ductal populations. The bottleneck score (Figure 6C) highlights this region with the highest intensity, consistent with the score capturing cells at the EMT–cell-cycle junction.

**Figure 6.**
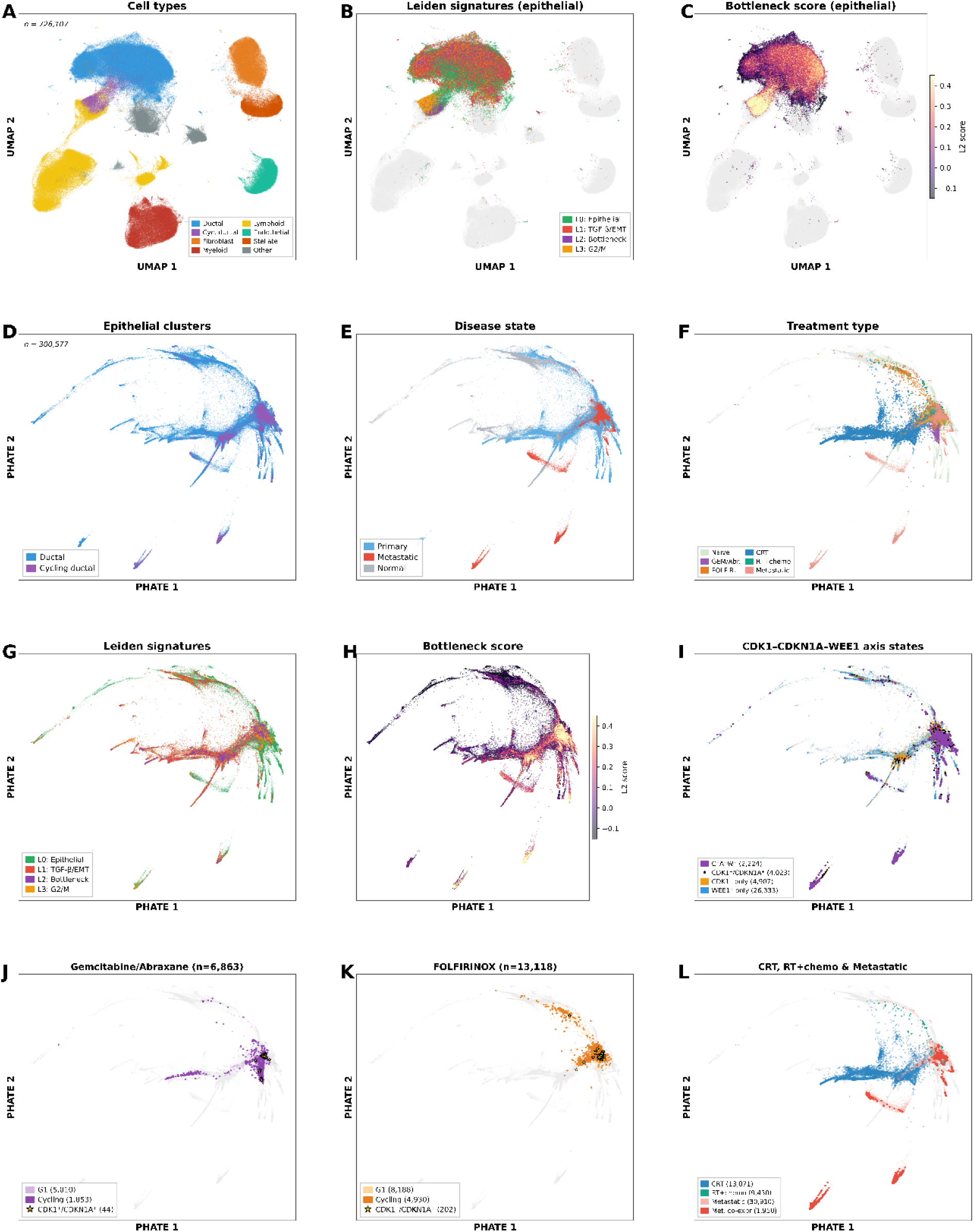
Multi-level mapping of CDK1–CDKN1A–WEE1 bottleneck signatures in the PDAC single-cell atlas. (A–C) Full atlas view (726,107 cells, 231 patients, 12 studies). (A) UMAP embedding colored by cell-type annotation, showing the major compartments: ductal (blue), cycling ductal (purple), fibroblast (orange), myeloid (red), lymphoid (yellow), endothelial (teal), stellate/pericyte (dark orange), and acinar/endocrine (grey). (B) Leiden signature assignment projected onto the full atlas (epithelial cells only; non-epithelial in light grey). Cells are colored by the dominant signature score among L0 (epithelial baseline, green), L1 (TGF-β/EMT, red), L2 (S-phase + EMT bottleneck, purple), and L3 (G2/M proliferative, orange). (C) Continuous Leiden 2 (bottleneck) score overlaid on epithelial cells (magma colormap). **(D–F)** Epithelial subset (300,577 cells) visualized by PHATE embedding, which preserves continuous trajectory structure. (D) Cluster annotation distinguishing ductal and cycling ductal populations. (E) Disease-state mapping: treatment-naïve primary tumors (blue), metastatic lesions (red), and normal/donor tissue (grey). (F) Treatment-type overlay showing the distribution of chemotherapy survivors: gemcitabine/abraxane (purple), FOLFIRINOX (orange), chemoradiation (blue), RT + chemotherapy (teal), and metastatic (pink). Naïve cells are shown in pale green as background. **(G–I)** Bottleneck signature and axis-state mapping on epithelial PHATE. (G) Dominant Leiden signature assignment. (H) Continuous Leiden 2 (bottleneck) score. (I) CDK1–CDKN1A–WEE1 combinatorial axis states: triple-positive cells (C^+^A^+^W^+^, n = 2,224; purple), CDK1^+^/CDKN1A^+^ co-expressing cells (black), CDK1^+^-only cells (orange), and WEE1^+^-only cells (blue). **(J–L)** Treatment-specific views. (J) Gemcitabine/abraxane survivors (n = 6,863) with G1-arrested cells in light purple and cycling cells in dark purple; CDK1^+^/CDKN1A^+^ co-expressing cells are highlighted as yellow stars. (K) FOLFIRINOX survivors (n = 13,118), same color scheme in orange tones. (L) Remaining treatment groups and metastatic cells: CRT (blue), RT + chemotherapy (teal), and metastatic cells (pink) with CDK1^+^/CDKN1A^+^ co-expressing metastatic cells in dark red.

Sub-setting to 300,577 epithelial cells and recomputing a PHATE embedding (Figure 6D–F) revealed the expected structure: ductal cells dominate the main body, with cycling ductal cells forming extended branches (Figure 6D). Disease-state mapping (Figure 6E) showed that metastatic cells (red) are enriched at the tips of these branches—the regions of highest proliferative activity—while normal cells (grey) concentrate in quiescent zones. Treatment-type overlay (Figure 6F) demonstrates that chemotherapy survivors from different regimens (gemcitabine/abraxane, purple; FOLFIRINOX, orange) occupy partially overlapping but distinct regions of the epithelial landscape.

Leiden signature mapping on the epithelial PHATE (Figure 6G) confirmed that the four signatures partition the epithelial compartment into biologically coherent domains: L0 (epithelial baseline, green) occupies the core, L1 (TGF-β/EMT, red) extends into mesenchymal-shifted branches, L3 (G2/M, orange) marks actively dividing cells, and L2 (bottleneck, purple) localizes specifically at the junction between L1 and L3 territories. The continuous bottleneck score (Figure 6H) forms a gradient peaking precisely at this junction. CDK1–CDKN1A–WEE1 axis states (Figure 6I) confirmed that triple-positive cells (C^+^A^+^W^+^, n = 2,224, purple) localize to the bottleneck region, while CDK1^+^-only cells (orange) spread along proliferative branches and WEE1^+^-only cells (blue) mark the quiescent compartment.

Treatment-specific views (Figure 6J–L) revealed striking differences in how chemotherapy survivors populate the epithelial landscape. Gemcitabine-treated cells (Figure 6J; n = 6,863) are predominantly G1-arrested (73%), with cycling survivors sparse and axis co-expressing cells (yellow stars) rare. FOLFIRINOX-treated cells (Figure 6K; n = 13,118) retain more cycling cells, with co-expressing cells visible in the bottleneck region. Metastatic cells (Figure 6L) show the broadest distribution, with CDK1^+^/CDKN1A^+^ co-expressing cells (dark red) concentrated at the proliferative tips—the most aggressive region of the landscape.

#### The bottleneck axis predicts metastatic association through additive component enrichment

We tested whether the three-component axis shows additive metastatic enrichment. Among CDK1^+^ cycling cells (n = 9,697), the proportion originating from metastatic lesions increased incrementally with each additional axis component (Figure 7A): CDK1^+^ alone, 29.3% metastatic; with co-expressed CDKN1A, 40.0% (+10.7 percentage points); with WEE1, 43.9% (+14.6%); with both CDKN1A and WEE1, 58.9% (+29.6%). Each component independently and additively contributes to metastatic enrichment, consistent with a three-part functional unit rather than coincidental co-detection.

**Figure 7.**
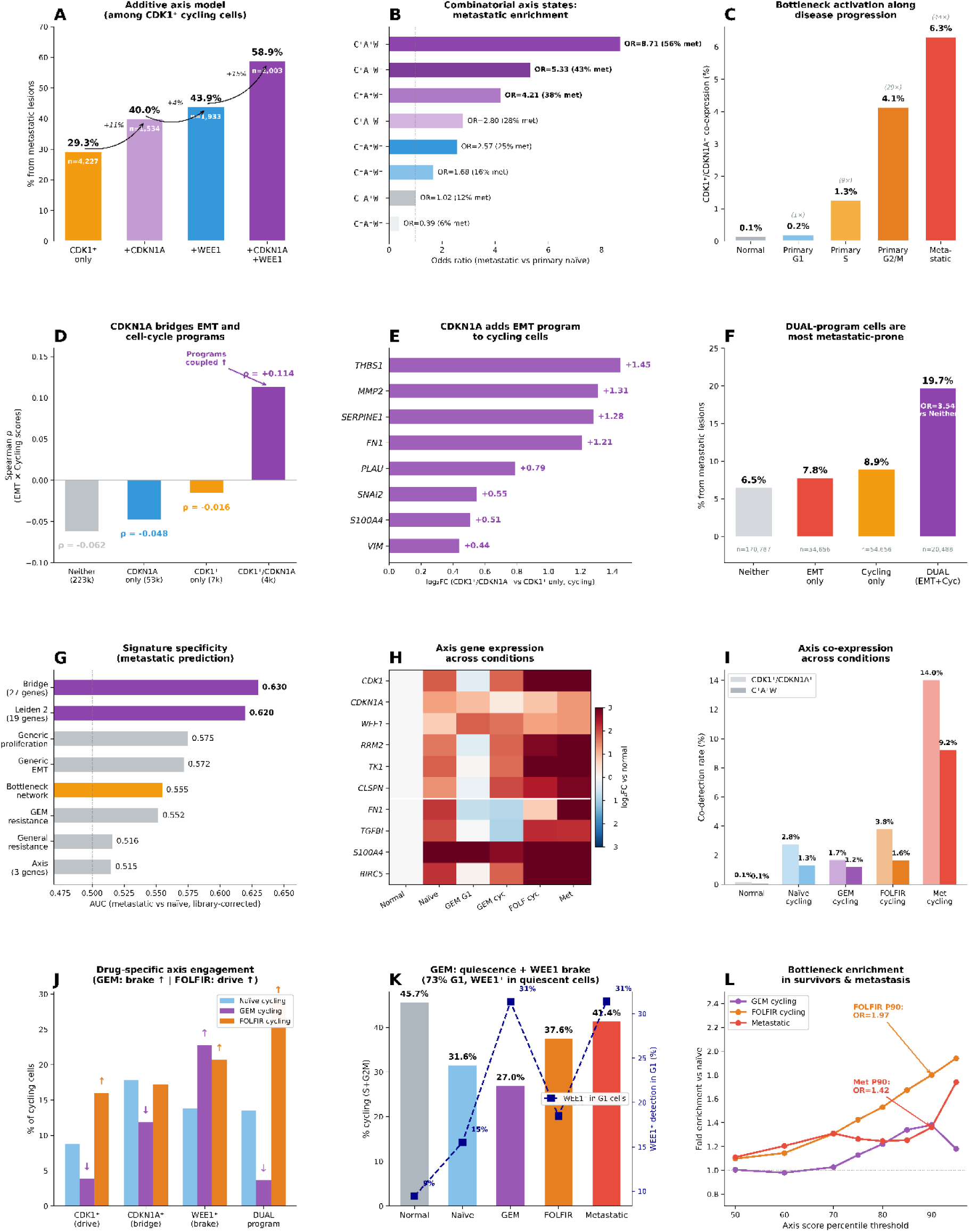
Quantitative validation of the CDK1–CDKN1A–WEE1 bottleneck axis: metastatic association, bridge mechanism, and treatment-specific patterns. (A–C) Metastatic association. (A) Additive axis model: among CDK1^+^ cycling cells (n = 9,697), the fraction originating from metastatic lesions increases with each additional axis component (CDK1^+^ only: 29.3%; +CDKN1A: 40.0%; +WEE1: 43.9%; +CDKN1A+WEE1: 58.9%). (B) Odds ratios for metastatic enrichment across all eight combinatorial states of CDK1/CDKN1A/WEE1 expression (Fisher’s exact test versus treatment-naïve primary; OR and percent metastatic shown for each state). Triple-positive cells show OR = 8.71 (56% metastatic); triple-negative cells are protective (OR = 0.39, 6.5% metastatic). (C) CDK1^+^/CDKN1A^+^ co-expression rate along disease progression, from normal epithelium (0.1%) through primary tumor cell-cycle phases (G1: 0.2%; S: 1.3%; G2/M: 4.1%) to metastatic cells (6.3%). Fold enrichment over normal is annotated above each bar. (D–F) Bridge mechanism. (D) Spearman correlation between aggregate EMT and cell-cycle program scores, stratified by CDK1/CDKN1A expression state. In non-expressing cells, programs are anti-correlated (ρ = −0.062); in CDK1^+^/CDKN1A^+^ cells, the correlation reverses to positive (ρ = +0.114, p = 5.1 × 10^−13^), demonstrating that co-expression specifically couples the two programs. (E) Log_2_ fold change of EMT genes in CDK1^+^/CDKN1A^+^ versus CDK1^+^-only cycling cells, showing that CDKN1A co-expression adds an EMT transcriptional program to actively cycling cells. (F) Fraction of cells from metastatic lesions across four program categories: neither EMT nor cycling (6.5%), EMT only (7.8%), cycling only (8.9%), and DUAL-program cells activating both (19.7%; OR = 3.54 versus neither). (G–I) Signature specificity. (G) AUC for discriminating metastatic versus treatment-naïve cells, corrected for library size, across multiple gene signatures. Bridge score (27 genes, AUC = 0.630) and Leiden 2 (19 genes, AUC = 0.620) outperform generic proliferation (0.575), generic EMT (0.572), and established resistance panels (0.515–0.555). Bottleneck network shown in orange. (H) Heatmap of log_2_ fold change for axis and associated genes across conditions relative to normal epithelium. White line separates cell-cycle genes (top) from EMT genes (bottom). (I) CDK1^+^/CDKN1A^+^ co-detection (light bars) and triple-positive C^+^A^+^W^+^ co-detection (dark bars) rates across conditions, showing progressive enrichment from normal (0.1%) to metastatic cycling cells (14.0% and 9.2%, respectively). (J–L) Treatment-specific patterns. (J) Prevalence of each axis component among cycling cells from treatment-naïve (blue), gemcitabine (purple), and FOLFIRINOX (orange) groups. Arrows indicate significant enrichment (↑) or depletion (↓) relative to naïve. Gemcitabine depletes CDK1^+^ (OR = 0.42) and DUAL-program cells (OR = 0.25), but enriches WEE1^+^ (OR = 1.84); FOLFIRINOX enriches all components. (K) Fraction of cycling cells (S+G2/M) by treatment group (bars), with WEE1^+^ detection rate among G1-arrested cells overlaid (blue dashed line, right axis). Gemcitabine survivors are most quiescent (27.0%) with highest WEE1 G1-detection (31%), consistent with WEE1-mediated checkpoint arrest. (L) Fold enrichment over treatment-naïve cells at increasing axis-score percentile thresholds for gemcitabine cycling survivors (purple), FOLFIRINOX cycling survivors (orange), and metastatic cells (red). Progressive enrichment at higher thresholds confirms dose-dependent bottleneck engagement.

Systematic analysis of all eight combinatorial states of the three-gene axis (Figure 7B) confirmed a hierarchy of metastatic association. The triple-positive state (C^+^A^+^W^+^) showed the strongest enrichment (OR = 8.71, 56.5% metastatic, Fisher’s exact test versus treatment-naïve primary), while the triple-negative state (C^−^A^−^W^−^) was protective (OR = 0.39, 6.5% metastatic). CDK1 positivity was the strongest single driver: all CDK1^+^ states exceeded OR > 2.8, whereas CDK1^−^ states without WEE1 showed minimal enrichment.

CDK1^+^/CDKN1A^+^ co-expression increased monotonically along disease progression (Figure 7C): 0.1% in normal epithelium, 0.2% in primary G1 cells, 1.3% in primary S-phase, 4.1% in primary G2/M, and 6.3% in metastatic cells—a 63-fold enrichment from normal to metastatic. This gradient parallels the expected trajectory from quiescence through cell-cycle entry to invasive capacity.

#### CDKN1A bridges EMT and cell-cycle programs specifically in co-expressing cells

A central prediction of the bottleneck model is that CDKN1A does not merely co-occur with CDK1 but functionally couples the EMT and cell-cycle modules. We tested this by computing the Spearman correlation between aggregate EMT and cell-cycle program scores across four mutually exclusive expression states (Figure 7D). In cells expressing neither gene (n = 223,000), EMT and cell-cycle scores were anti-correlated (ρ = −0.062), reflecting the established antagonism between these programs. This anti-correlation persisted in CDKN1A^+^-only cells (ρ = −0.048) and diminished but remained negative in CDK1^+^-only cells (ρ = −0.016). Strikingly, in CDK1^+^/CDKN1A^+^ co-expressing cells, the correlation reversed to positive (ρ = +0.114, p = 5.1 × 10^−13^), indicating that CDKN1A co-expression specifically enables the simultaneous activation of both programs—the defining feature of the bottleneck state.

To identify the molecular basis of this coupling, we compared EMT gene expression between CDK1^+^/CDKN1A^+^ co-expressing cells and CDK1^+^-only cells, restricting to cycling cells to control for cell-cycle stage (Figure 7E). Co-expression of CDKN1A was associated with marked upregulation of EMT effectors: *THBS1* (log_2_FC = +1.45), *MMP2* (+1.31), *SERPINE1* (+1.28), *FN1* (+1.21), *PLAU* (+0.79), *SNAI2* (+0.55), and *VIM* (+0.44; all p < 10^−76^). Thus, CDKN1A adds an EMT transcriptional program on top of an already active cell cycle—precisely the bridge function predicted by the bottleneck model.

Cells activating both EMT and cell-cycle programs simultaneously (DUAL-program cells, defined as scoring above the 75th percentile for both L1 and L3 signatures) constituted the most metastatic-prone population in the atlas (Figure 7F): 19.7% of DUAL cells originated from metastatic lesions, compared with 8.9% of cycling-only cells, 7.8% of EMT-only cells, and 6.5% of cells in neither program (OR = 3.54 versus neither, p < 10^−100^). CDK1^+^/CDKN1A^+^ co-expressing cells were 8.62-fold enriched for DUAL-program membership (37.1% versus 5.5% in neither-expressing cells), confirming that the bottleneck axis marks cells that have successfully coupled both modules.

#### The bottleneck signature outperforms generic and resistance-associated gene sets

To assess whether the bottleneck captures biology beyond generic proliferation or EMT, we compared multiple gene signatures for their ability to discriminate metastatic from treatment-naïve cells (Figure 7G). All scores were corrected for library size to remove technical confounders. The 27-gene bridge signature (genes uniquely elevated in DUAL-program cells) achieved the highest AUC of 0.630, followed by the 19-gene Leiden 2 signature (AUC = 0.620). Generic proliferation (AUC = 0.575), generic EMT (AUC = 0.572), and established drug-resistance gene panels (GEM resistance AUC = 0.552, general resistance AUC = 0.516) performed substantially worse. The three-gene axis alone (CDK1/CDKN1A/WEE1, AUC = 0.515) was limited as a continuous score due to single-cell dropout, but its binary co-detection pattern yielded the strongest categorical association (OR = 8.71).

Gene-level expression analysis across disease and treatment conditions (Figure 7H) confirmed that the bottleneck axis genes are upregulated in metastatic cells relative to normal epithelium, with cell-cycle components (*CDK1*, *RRM2*, *TK1*, *CLSPN*) and EMT genes (*FN1*, *TGFBI*, *S100A4*) co-elevated in metastatic tissue. Notably, gemcitabine G1-arrested survivors showed selective *WEE1* upregulation without *CDK1* elevation, consistent with checkpoint-mediated arrest. FOLFIRINOX cycling survivors showed broad upregulation of both cell-cycle and EMT axis genes.

Axis co-expression rates across conditions (Figure 7I) revealed that CDK1^+^/CDKN1A^+^ co-detection was highest in metastatic cycling cells (14.0%) and FOLFIRINOX cycling survivors (3.8%), compared with 2.8% in naïve cycling cells and 0.1% in normal epithelium. Triple-positive co-detection (C^+^A^+^W^+^) followed the same hierarchy: 9.2% metastatic, 1.6% FOLFIRINOX, 1.2% GEM, and 1.3% naïve cycling.

#### Treatment survivors show drug-specific axis engagement consistent with distinct survival strategies

Comparing axis component prevalence between treatment-naïve and chemotherapy-surviving cycling cells (Figure 7J) revealed opposing patterns for gemcitabine and FOLFIRINOX. Gemcitabine cycling survivors were depleted for CDK1^+^ cells (3.9% versus 8.8% naïve, OR = 0.42, p < 10^−16^) but enriched for WEE1^+^ (22.8% versus 13.8%, OR = 1.84, p < 10^−24^)—a pattern consistent with WEE1-mediated G2/M checkpoint arrest as the dominant survival mechanism. The DUAL program was strongly depleted in gemcitabine survivors (3.7% versus 13.5%, OR = 0.25), suggesting that gemcitabine effectively eliminates cells in the active bottleneck state. FOLFIRINOX survivors, by contrast, showed enrichment for CDK1^+^ (16.0%, OR = 1.96), DUAL program (28.9%, OR = 2.61), and WEE1^+^ (20.7%, OR = 1.63)—consistent with continued cell-cycle transit through the bottleneck despite treatment.

The quiescence pattern (Figure 7K) reinforced this interpretation. Gemcitabine survivors were the most quiescent population (27.0% cycling, compared with 31.6% naïve, OR = 0.80), with their G1-arrested cells showing elevated WEE1 detection (31% versus 16% in naïve G1), consistent with active checkpoint engagement. Metastatic cells showed the opposite trend: 41.4% cycling, indicating that cells reaching metastatic sites maintain high proliferative capacity.

Axis score percentile enrichment (Figure 7L) provided a dose-response view of bottleneck engagement across conditions. At the 90th percentile of the axis score, FOLFIRINOX cycling survivors showed 1.97-fold enrichment over naïve cells, metastatic cells 1.42-fold, and gemcitabine survivors 1.44-fold. The progressive enrichment at higher percentile thresholds confirms that the most extreme bottleneck activation is selectively retained in both treatment survivors and metastatic cells.

We note that each treatment arm in this atlas derives from a single study with limited patient numbers, and pre-treatment samples from the same patients are not available. We therefore cannot formally distinguish drug selection from patient-intrinsic biology. The observed patterns are consistent with drug-specific survival strategies but require validation in matched pre- and post-treatment cohorts.

## Discussion

In this study, we applied dynamic single-cell trajectory analysis combined with network state-transition inference to 3D PANC-1 tissue models to characterize regulatory mechanisms underlying gemcitabine resistance under conditions that approximate the clinical complexity of PDAC. Our matrix-based models show pronounced chemoresistance that is further amplified by TGF-β1 stimulation, recapitulating the interplay between EMT and drug resistance observed in patient tumors. By reconstructing developmental paths and state transitions over pseudo-time—rather than relying on static expression snapshots—we resolved attractor states and identified candidate regulators of resistance. Superimposing network-based logic from protein–protein interactions onto these trajectories added mechanistic context, highlighting not only correlated expression changes but candidate upstream drivers and their temporal sequence. External validation in a comprehensive PDAC single-cell atlas (726,107 cells, 231 patients, 12 studies) confirmed that the central regulatory architecture identified in vitro—the CDK1–CDKN1A–WEE1 bottleneck—is a recurrent feature of human PDAC biology associated with metastatic propensity, treatment-specific survival strategies, and the most aggressive disease phenotypes.

### Single-cell expression landscape and state transitions in 3D PANC tissue models

Single-cell data reveal that individual cells within a tissue exhibit highly diverse expression profiles, reflecting their inherently stochastic and dynamic nature (Wolfsberger et al., 2023; Wang et al., 2024). These states typically cluster around a limited number of attractor states—such as specific cell-cycle phases—and transitions between attractors require substantial changes in gene expression. In our analysis of PANC cells across four conditions (CTR, GEM, TGF-β1, GEM+TGF-β1), we observe high heterogeneity and discrete Leiden clusters corresponding to major cell-cycle phases (Fig. 2). Cells transitioning to S and G2/M show dramatic expression changes—including alterations in spindle assembly and E2F targets—indicating that specific drivers such as CDK1 and the TPX2–AURKA axis propel cells from quiescence into active cycling. These dynamics indicate that PANC(CTRL) cultures are actively cycling, beginning in Leiden 0 (quiescent G1) and progressing through Leiden 3 into S phase and ultimately G2/M, with two distinct trajectories: one consolidating S-phase arrest and the other transitioning through G2/M (Figs. S4A–D).

### Cell-cycle state transitions and intrinsic gemcitabine resistance

Our analysis indicates that Leiden 3—characterized by high expression of cell-cycle regulators (Mitotic spindle, mTORC1, E2F targets, G2–M checkpoint pathways)—is enriched for GEM-surviving cells. These cells are predominantly in late S/early G2 phase, suggesting S-phase locking. Key regulators including BIRC5, TOP2A, TPX2, AURKA, CENPF, RRM2, CDK1, CDKN1A, and KIF family genes appear to enable circumvention of GEM-mediated replication stress.

These findings are consistent with the established mechanism of GEM, a nucleoside analog that disrupts DNA synthesis and inhibits ribonucleotide reductase, thereby inducing S-phase arrest (Shewach & Lawrence, 1996). Cells arrested in S phase activate the ATR/CHK1 checkpoint axis, with many undergoing apoptosis and only a few progressing to G2/M (Shewach & Lawrence, 1996; Wang et al., 2009). Resistance-associated pathways include AURKA/TPX2, which have been linked to gemcitabine resistance by shortening S-phase duration (Marumoto et al., 2005; Lens et al., 2010; Guenther et al., 2023); pro-survival signaling from BIRC5 (Li, 2007) and mTORC1 activation (Son et al., 2024; Kagawa et al., 2012; Chawsheen et al., 2021); CDK1 dysregulation enabling bypass of G2–M checkpoints (Wang et al., 2023; Wijnen et al., 2021); and RRM2-driven nucleotide pool replenishment supporting DNA repair (Davidson et al., 2004; Jordheim & Dumontet, 2007; Zhan et al., 2021). These findings suggest that specific subpopulations within 3D tissues may possess intrinsic gemcitabine resistance through S-phase arrest or bypass of the S→G2 transition.

### TGF-β1-induced EMT state transitions and resistance

Beyond cell-cycle–driven transitions, TGF-β1 stimulation introduced an additional attractor in our experiments. TGF-β1 signaling induces EMT through both canonical SMAD-mediated transcription and non-canonical pathways including PI3K/AKT, KRAS/MAPK, and mTOR (Chen et al., 2018). TGF-β1 appears to push cells from multiple starting states toward a new differentiation trajectory marked by EMT, UV response, and KRAS-associated pathways. This effect manifests in parallel across quiescent (G1) and S-enriched root states, suggesting that TGF-β1 overrides inherent attractors and reorients cells toward an EMT-like phenotype. Temporal clustering highlights key EMT drivers (COL5A1, THBS1, FN1, COL1A1, MMP2, TGFBI) with concurrent downregulation of epithelial markers (KRT18, KRT19). Notably, many late-induced genes have only begun to increase, suggesting an incomplete shift toward a stable EMT attractor that may require additional time or signals.

Several late-responding genes are strongly linked to poor prognosis in pancreatic cancer. S100A2 is a potent EMT driver implicated in gemcitabine resistance (Chen et al., 2021; Mahon et al., 2007). FN1 is a key metastatic marker similarly implicated in resistance (Xavier et al., 2021). MBOAT2 has been identified as an unfavorable biomarker (Li et al., 2022), and PGM2L1, a glucose 1,6-bisphosphate synthase, has emerged as a novel prognostic biomarker in cholangiocarcinoma with implications for chemoresistance and immune evasion (Wu et al., 2025). These observations align with the broader literature linking EMT to both metastasis and drug resistance in pancreatic cancer (Du et al., 2024; El Amrani et al., 2019; Weadick et al., 2021; Debaugnies et al., 2023). Our data suggest that TGF-β1-induced EMT contributes to chemoresistance independently, potentially explaining why Leiden 1 is enriched with gemcitabine survivors.

### Combined G1→S and TGF-β1/EMT transitions synergistically enhance resistance

The combination of G1→S progression and TGF-β1 stimulation produces a synergistic phenotype: EMT-positive cells arrested in S phase after gemcitabine treatment. Genes characterizing this hybrid state include TYMS, UBE2T, ATAD2, TK1, MYBL2, RRM2, HMGB1, MELK, CENPW, and DTL. Among these, MYBL2 is a central transcription factor associated with chemoresistance and poor clinical outcomes (Musa et al., 2017); HELLS is a chromatin remodeler linked to accelerated tumor growth and reduced therapy responsiveness (Hou et al., 2020; Zocchi et al., 2020); and DTL, which closely tracks gemcitabine-survivor enrichment (Fig. 4D), is a key hypoxia target regulated by HIF-1α (Lovisa et al., 2015; Akhmetkaliyev et al., 2023). These observations suggest that intrinsic resistance mechanisms—such as rapid S-phase entry, efficient DNA repair, and nucleotide pool maintenance—can be reinforced by EMT-driven S-phase arrest. Importantly, TGF-β1-stimulated EMT can arrest the cell cycle while concurrently promoting motility (Song, 2007; Lovisa et al., 2015; Akhmetkaliyev et al., 2023), suggesting that Leiden-2-like S-phase-EMT cells constitute a high-risk subpopulation with both pronounced chemoresistance and metastatic potential.

### Network analysis identifies a CDK1–CDKN1A bottleneck as the emergent decision point

To understand why resistance emerges specifically from the combination of cell-cycle progression and EMT, we tracked dynamic network changes across cell-state trajectories. Several genes showed both high network reach and strong expression impact in the combined state, including DTL, E2F1, CLSPN, CCNA2, BRIP1, RRM2, and MYBL2—factors linked to replication stress and cell-cycle regulation that were especially expressed in Leiden cluster 2. Mechanistically, RRM1/RRM2 buffer nucleotide depletion (Davidson et al., 2004; Jordheim & Dumontet, 2007); DTL activates the CRL4Cdt2 E3 ubiquitin ligase promoting p21 degradation during S phase (Abbas & Dutta, 2011); MYBL2 and E2F1 drive the transcriptional program for S→G2 progression; and CLSPN, RAD51, and BRIP1 safeguard replication fork stability (Petermann et al., 2010).

Analysis of betweenness centrality changes along treatment trajectories revealed a striking emergent bottleneck centered on CDK1 and CDKN1A (p21), uniquely triggered when G1→S progression and EMT co-occurred. Either pathway alone modestly increased CDK1 centrality, but their combination caused a sharp spike, indicating that CDK1 became a critical bridge between major network modules. Shortest-path mapping identified SMAD3 and CDK4 as primary upstream conduits. SMAD3 drives TGF-β1-dependent p21 activation, while CDK4 promotes G1 progression via RB phosphorylation and E2F activation (Wijnen et al., 2021). An unbiased random knockout screen confirmed that CDKN1A had the most substantial impact on CDK1 centrality. CDK1 expression was primarily driven by S-phase progression, whereas CDKN1A induction was driven by TGF-β1 signaling. The combined state thus produced a “poised but arrested” condition—cells primed for division yet held in S-phase persistence.

This interpretation aligns with evidence that EMT programs promote chemoresistance by enhancing replication-stress tolerance and activating the DNA damage response (Debaugnies et al., 2023; Zheng et al., 2015; Schuhwerk & Brabletz, 2023). The marked elevation of p21 emerges as a central survival mechanism, enabling cells to persist despite DNA damage (Gartel & Tyner, 2002). Mechanistically, TGF-β1 can induce p21 transcription in a p53-independent manner (Datto et al., 1995), and once elevated, p21 directly inhibits CDK1, blocking the S→M transition under genotoxic stress. Together, these findings support our inference that CDK1 and CDKN1A function as a critical bottleneck: CDKN1A enforces arrest while CDK1 stocks the mitotic machinery, fostering an S-phase persistence state where cells survive gemcitabine by arresting before mitosis—buoyed by fork protection, HR repair, and robust nucleotide supply—poised to resume proliferation post-stress.

### Experimental context: proliferation in 3D tissue models

We observed that GEM treatment leads in our tissue-like conditions to a higher Ki-67 proliferation index, in contrast to the viability reduction typical of conventional 2D models with artificially high proliferation indices (Namima et al., 2020). Our matrix-based PDAC tissue model has a proliferation index of approximately 35%, better reflecting the clinically relevant mean of 26% Ki67-positive cells in PDAC patients (Striefler et al., 2016). Notably, in that same study of 162 PDAC patients, GEM-treated tumors with lower Ki67 indices showed longer PFS and OS. TGF-β1 acts differently on cancer cells than on healthy cells: while it typically induces G1 arrest in healthy cells, recent evidence indicates that TGF-β1-induced EMT can be accompanied by increased proliferation in cancer, particularly when interconnected with DNA damage response and cell-cycle progression (Schuhwerk & Brabletz, 2023). Clinical trials targeting TGF-β1 (galunisertib; Melisi et al., 2018) or EMT (trabectedin; Ko et al., 2020) have shown only modest survival benefits in PDAC, underscoring the need for better patient stratification with novel targets.

### External validation contextualizes the bottleneck as a clinically relevant regulatory state

Our tissue-culture model provides high-resolution characterization of a controlled perturbation, allowing identification of the CDK1–CDKN1A–WEE1 bottleneck as a structural feature of the gene-regulatory network. The PDAC atlas analysis demonstrates that this bottleneck reflects a biologically and clinically meaningful state in human tumors. First, triple-positive (CDK1+/CDKN1A+/WEE1+) cells localize to a specific region of the epithelial landscape—the junction between proliferative and mesenchymal branches—rather than being distributed randomly, arguing against technical artifacts. Second, the additive architecture (each component independently contributing to metastatic risk, from 29% with CDK1+ alone to 59% with all three) suggests a modular functional unit. The 8.71-fold enrichment for metastatic origin among triple-positive cells exceeds generic proliferation, EMT, and established drug-resistance gene panels. Third, the correlation-reversal analysis provides molecular evidence for the bridging function of CDKN1A: the EMT–cell-cycle anti-correlation (ρ = −0.062) reverses to a positive coupling (ρ = +0.114) specifically in CDK1+/CDKN1A+ co-expressing cells, consistent with our network model. The resulting DUAL-program cells constitute the most metastatic-prone epithelial population in the atlas (19.7% metastatic, 77% basal-like).

Treatment data suggest the bottleneck axis may have clinical relevance beyond metastatic prediction. Gemcitabine survivors show a ‘brake-only’ pattern: depletion of CDK1+ cells and enrichment of WEE1+ quiescent cells, consistent with checkpoint-mediated arrest as a survival strategy. FOLFIRINOX survivors, by contrast, maintain active cycling through the bottleneck with enrichment of the full DUAL program. These observations are independently supported by Werba et al. (2023), who found that chemotherapy-treated PDAC cells were enriched for EMT programs compared to treatment-naïve tumors, consistent with our model in which chemotherapy selects for cells engaging the EMT–cell-cycle bottleneck. The controlled tissue model and atlas analysis are complementary: the tissue model provides mechanistic resolution to identify and characterize the bottleneck’s regulatory logic, while the atlas provides statistical power to establish that this architecture is a recurrent feature of human PDAC biology.

### Treatment implications and future directions

The CDK1/CDKN1A bottleneck topology highlights several promising intervention points in clinical development. The most direct vulnerability is the ATR–CHK1–WEE1 checkpoint axis, which stabilizes replication forks and maintains CDK1 in an inactive, p21-bound state. Inhibiting these kinases with gemcitabine can collapse S-phase persistence, forcing premature mitotic entry (Parsels et al., 2013; Karnitz & Zou, 2015). Clinical translation is underway: the WEE1 inhibitor adavosertib combined with gemcitabine and radiation achieved a median OS of 21.7 months in locally advanced PDAC (Cuneo et al., 2019), while the CHK1 inhibitor SRA737 with gemcitabine showed a favorable tolerability profile (Jones et al., 2023), and LY2880070 with gemcitabine highlighted the need for better stratification biomarkers (Huffman et al., 2023). Our bottleneck model may provide such a stratification tool, as the atlas reveals two distinct survival strategies: gemcitabine survivors adopt a ‘brake-only’ pattern amenable to WEE1 inhibition, while FOLFIRINOX survivors maintain active cycling through the full axis, suggesting benefit from combined ATR–CHK1–WEE1 targeting.

Upstream, CDK4/6 inhibition can dampen the E2F/MYBL2 program that elevates CDK1, though monotherapy shows limited efficacy due to compensatory pathways and can paradoxically induce TGF-β/SMAD-dependent EMT (Saqub et al., 2023; Salvador-Barbero et al., 2020; Wang et al., 2025). The principle that durable PDAC therapy requires simultaneous multi-node disruption is illustrated by Liaki et al. (2025), whose triple combination of RAS(ON), EGFR, and STAT3 inhibitors achieved complete PDAC regression without resistance. Notably, STAT3 mediates TGF-β-induced EMT and can activate p21 transcription (Bromberg et al., 1999), providing a molecular link to our bottleneck. These converging findings suggest that combining checkpoint inhibitors with agents targeting EMT and KRAS signaling axes may be necessary to dismantle the network architecture sustaining resistance. Future work should evaluate whether bottleneck signature scores can serve as predictive biomarkers for patient stratification in checkpoint inhibitor trials.

### Conclusion

Our work introduces a physiologically relevant 3D PDAC model that enables single-cell mapping of drug-induced resistance trajectories. We identify a novel ‘S-phase persistence’ hybrid state where proliferative drive and TGF-β1-induced restraint converge at the CDK1–CDKN1A–WEE1 bottleneck, generating a chemo-tolerant subpopulation likely underlying relapse and minimal residual disease. Cross-examination in a PDAC atlas comprising 726,107 cells from 231 patients across 12 studies confirms this bottleneck as a recurrent feature of human PDAC biology: triple-positive cells show 8.7-fold enrichment for metastatic origin, CDKN1A co-expression reverses the normal EMT–cell-cycle anti-correlation, and GEM survivors show a WEE1-dominant checkpoint arrest pattern distinct from the full DUAL program enrichment seen in FOLFIRINOX survivors — suggesting the axis captures a general chemoresistance mechanism. These findings point to actionable vulnerabilities in ATR–CHK1–WEE1 and TGF-β1/SMAD3 signaling and suggest that bottleneck engagement patterns could stratify patients for combination therapies — consistent with the emerging principle that durable PDAC responses require simultaneous disruption of multiple signaling nodes (Liaki et al., 2025). Together, these findings establish both a biological model and a methodological framework applicable to other cancer contexts where single-cell resolution is available.

## Credit author statement

MA: **Conc**-**Meth**-**Soft**-**Val**-**Form**-**Invest**-Res-**DataC**-**WriteO**-**WriteRE**-**Vis**-Sup-Adm-Fund

JGNP: Conc-**Meth**-Soft-**Val**-**Form**-**Invest**-Res-**DataC**-**WriteO**-WriteRE-**Vis**-Sup-Adm-Fund

TD: **Conc**-Meth-Soft-Val-**Form-Invest**-**Res**-DataC-**WriteO**-**WriteRE**-Vis-**Sup**-**Adm-Fund**

GD: **Conc**-Meth-Soft-Val-Form-**Invest**-**Res**-DataC-**WriteO**-**WriteRE**-Vis-**Sup**-**Adm-Fund**

JB: **Conc**-**Meth**-**Soft**-**Val**-**Form**-**Invest**-Res-**DataC**-**WriteO**-**WriteRE**-**Vis**-**Sup**-**Adm**-Fund

## Acknowledgements

The authors acknowledge the use of ChatGPT (OpenAI, San Francisco, USA) and Grammarly for language polishing, text condensation, and minor code optimization. All content, including the conceptual design, analysis, and interpretation, was created, verified, and approved by the authors. The authors take full responsibility for the integrity and accuracy of the manuscript.

## Availability of data and materials

The raw single-cell RNA sequencing data generated in this study have been deposited in ArrayExpress (accession number pending final metadata curation; https://www.ebi.ac.uk/biostudies/arrayexpress). The processed annotated dataset will be made available via the EMBL-EBI Single Cell Expression Atlas upon accession approval. The PDAC single-cell atlas used for external contextualization is publicly available from Zenodo (accession 14199536; Loveless et al., 2025). Analysis code and conda environment YAML files for full reproducibility are available at https://github.com/Almasimaryam/scRNAseq-pipeline-Py.git.

## Additional files

Supplementary Tables (including Table S1 and S2). Supplementary data (including Figures S1-S7, additional material and methods, and additional discussion)

## Ethics approval and consent to participate

All procedures involving porcine tissue were conducted in accordance with institutional and national guidelines for animal welfare. Collection of porcine intestinal tissue was performed under the approval of the Ethics Committee of the District of Unterfranken, Würzburg, Germany, approval number #2532-2-12. No human participants were involved in this study.

